# Chloroplast-encoded small subunit extensions reshape the Chlamydomonas chlororibosome

**DOI:** 10.64898/2026.02.07.704542

**Authors:** Florent Waltz, Philippe A. Lehner, Philippe Van der Stappen, Lukas Kater, Stefan Pfeffer, Benjamin D. Engel

## Abstract

Chloroplast ribosomes (chlororibosomes) synthesize the core protein components of the photosynthetic apparatus, yet their structural diversity outside flowering plants remains largely unexplored. Here, we combine in situ cryo-electron tomography (cryo-ET) with single-particle cryo-electron microscopy (cryo-EM) to determine the structure of the chlororibosome from the unicellular green alga *Chlamydomonas reinhardtii*. Subtomogram averaging of chlororibosomes in their native environment, resolved to ∼5 Å resolution and in distinct translational states, reveals particles both free in the stroma and loosely tethered to thylakoid membranes. These *in situ* reconstructions uncover an additional “arm” domain on the small subunit. High-resolution single-particle reconstruction of isolated chlororibosomes to ∼2.5 Å, in states bound either to the inhibitory translation factor pY or to a nascent chain-linked P-site tRNA, reveals that this domain is built primarily from extensive chloroplast-encoded insertions and extensions of conserved small subunit proteins, supported by chlororibosome-specific ribosomal proteins. The arm domain is located around the mRNA entry and exit channels, suggesting a role in stabilizing the mRNA trajectory through the small subunit and organizing chloroplast polysomes. Together, these data reveal unexpected structural variation of algal chlororibosomes and suggest that chloroplast translation has diversified substantially even among relatively closely related photosynthetic lineages.

## Introduction

Chloroplasts are key organelles of photosynthetic organisms, responsible for oxygenic photosynthesis. Their origins trace back to an ancient endosymbiotic event between a eukaryotic host cell and a cyanobacterial ancestor^1^. This evolutionary merger had a profound impact on our planet, as plants and microalgae constitute the base of most food chains, producing many molecules of interest for human health as well as being major players in carbon sequestration^2–4^. Chloroplasts have also emerged as attractive “green factories” for high-level expression of heterologous proteins and resistance traits^5–7^.

Chloroplasts contain their own ribosomes, chlororibosomes, which were inherited from cyanobacteria and are responsible for synthesizing ∼75 chloroplast-encoded proteins out of ∼1,000 proteins in the organelle^8–10^. Despite their central role in assembling and maintaining the photosynthetic apparatus, chlororibosomes remain much less structurally characterized than bacterial or cytosolic ribosomes. In contrast, mitochondrial ribosomes have been thoroughly studied and exhibit significant structural and compositional diversity among eukaryotes^11–13^. While the common consensus is that chlororibosomes are less diverse and more bacteria-like compared to mitoribosomes, our structural knowledge is solely based on chlororibosomes from the flowering plant *Spinacia oleracea*, commonly known as spinach^14–17^. However, chloroplasts are distributed across a broad phylogenetic spectrum^18^, making it likely that chlororibosomes vary in composition, architecture, and interactions with regulatory and assembly factors that control chloroplast biogenesis and photosynthetic capacity^19,20^. Defining this diversity is essential for understanding how chloroplast translation is tuned to support photosynthesis.

## Results

### The *C. reinhardtii* chlororibosome harbors an additional domain on its small subunit

To probe chlororibosome diversity beyond flowering plants, we worked with *Chlamydomonas reinhardtii*, a unicellular green alga that serves as a valuable model to study chloroplast biology due to its genetic tractability and established experimental techniques^21–23^. To image chlororibosomes in their most native state, we leveraged the publicly available Chlamydomonas cryo-ET dataset, EMPIAR-11830^24^. We selected 129 chloroplast-containing tomograms to perform automated chlororibosome detection using template matching and subsequent subtomogram averaging (Supplementary Fig. 1). Chlororibosomes were detected in the stroma, both as free and in association with polysomes, as well as bound to thylakoid membranes (Fig. 1). This observation indicates that chlororibosomes are actively translating both soluble stroma proteins and membrane proteins for the photosynthetic machinery. Interestingly, we also observed large polysomes organized at the membrane, often circular, a structural feature that was first described by Chua et al. in 1976^25^ (Fig. 1c).

**Figure 1:**
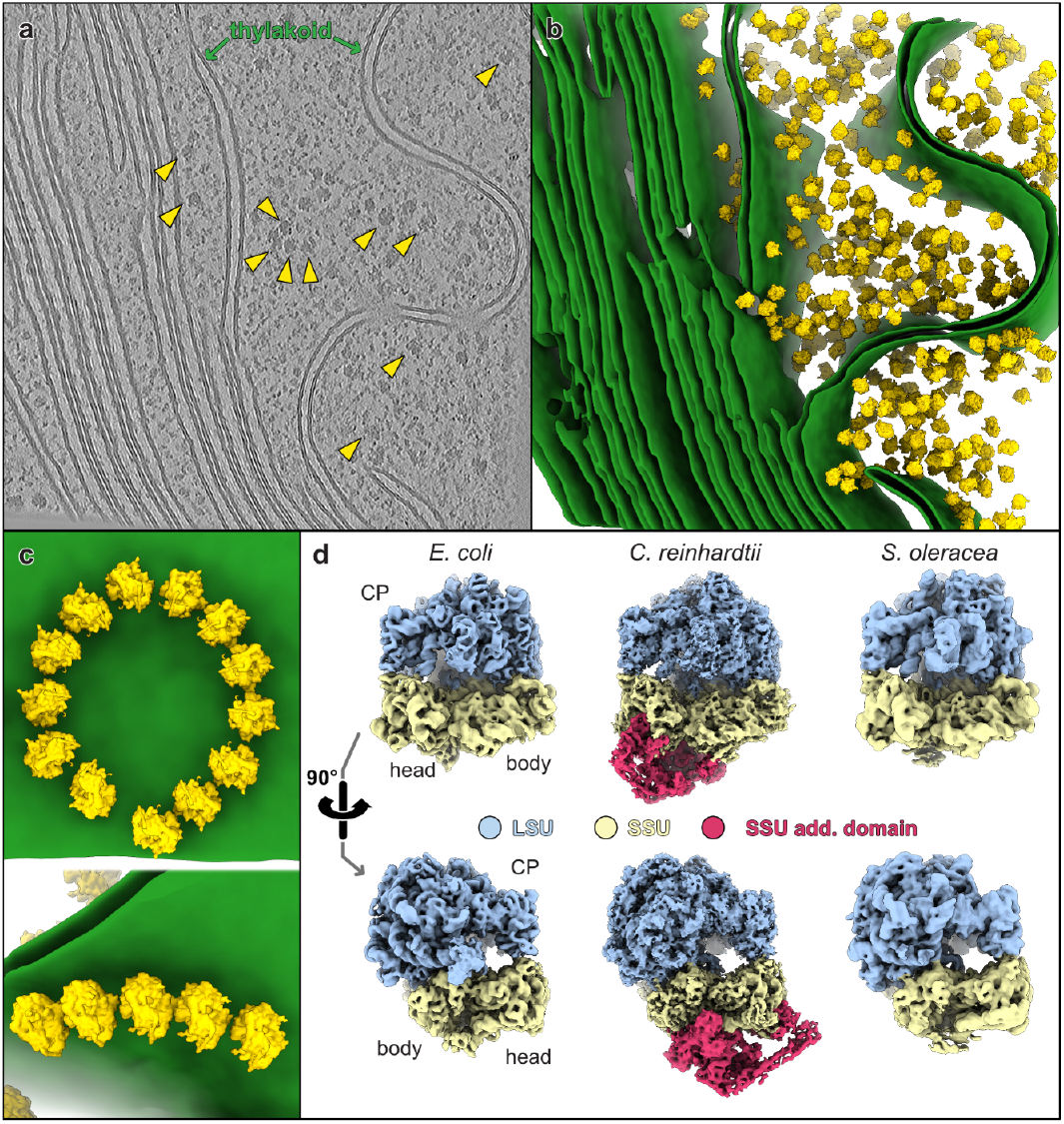
In-cell structure and organization of Chlamydomonas chlororibosomes. **a**, Slice of a tomogram depicting a chloroplast within a native *C. reinhardtii* cell. Yellow arrowheads point to chlororibosomes. **b**, Corresponding segmentation of the thylakoid membranes (green) with STA structures of the chlororibosomes (yellow) mapped back into their respective particle positions. **c**, Close up views of membrane-bound chloroplast polysomes. **d**, Subtomogram average of the *C. reinhardtii* chlororibosome, highlighting the large additional “arm” domain compared with the *E. coli* bacterial ribosome and the *S. oleracea* chlororibosome.

Subtomogram averaging (STA) of the chlororibosomes reached ∼5 Å resolution, revealing an unexpected architectural feature: a substantial additional domain on the small subunit. Since this domain roots from the body of the ribosome and extends toward the head, we name it hereafter the small subunit (SSU) arm domain (Fig. 1d). Comparison with bacterial ribosome and the spinach chlororibosome structures shows that this SSU domain is absent from previously characterized chlororibosomes, suggesting that it represents a species-specific adaptation in Chlamydomonas. The base of the SSU arm is located around the mRNA entry and exit channels, hinting that this additional domain could be made either from new r-proteins interacting there or from extensions of these conserved r-proteins.

### Thylakoid-associated chlororibosomes are loosely tethered yet actively translating *in situ*

Subtomogram classification revealed that only a fraction of chlororibosomes are positioned adjacent to thylakoid membranes (∼31%), whereas the majority remain free in the stroma (Fig. 2a). Thylakoid-associated ribosomes are relatively distant from the membrane surface compared to cytosolic or mitochondrial ribosomes (∼40 Å from the peptide exit channel to the membrane). In the average, only a weak, thin density bridges the ribosomal exit region (near rRNA helix H59) with the membrane (Supplementary Fig. 2a). This suggests that the tether, likely comprising the nascent polypeptide and/or components of a thylakoid protein translocase, is highly flexible or compositionally heterogeneous, allowing substantial lateral and angular motion of the ribosome relative to the membrane (Fig. 2b). These observations support a model in which chlororibosomes engage the thylakoid translocon in a dynamic manner, in contrast to cytosolic ER-bound ribosomes and mitochondrial ribosomes, which are more rigidly docked (Supplementary Fig. 2a). This is confirmed by the range of chlororibosome orientations observed relative to the thylakoid membrane, spanning 20° in the most extreme case (Fig. 2b). On the ribosome side, the area surrounding the peptide exit channel is particularly bacterial-like and does not show specific domains or remodeling that would support a specialized translocon interaction (Supplementary Fig. 2b). Consistently, bL23c is more similar to its bacterial counterpart than its spinach counterpart, arguing against a possible alternative exit channel in Chlamydomonas chlororibosomes^15^.

**Figure 2:**
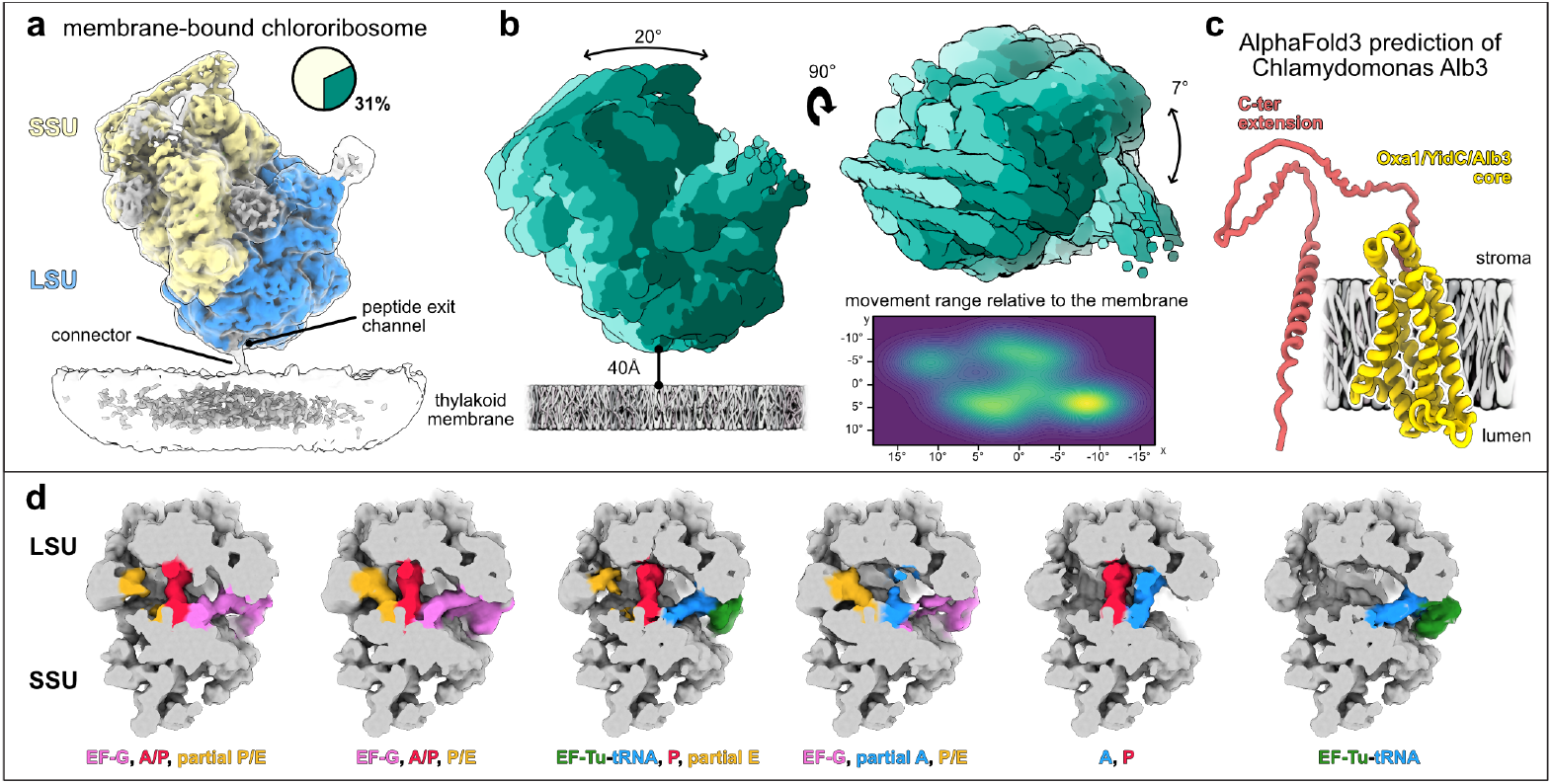
Membrane associated chlororibosomes and translation state. **a**, Subtomogram average of thylakoid-bound chlororibosomes, representing 31% of the particles. The thin proteinaceous connector adjacent to the peptide exit channel contacting the thylakoid membrane is shown. **b**, Analysis of chlororibosome movement relative to the membrane. The different orientations relative to the membrane are overlayed with different shades of teal, from the two different views. The heat map shows the projected movement range of the ribosome relative to the membrane plane. **c**, AlphaFold3 prediction of *C. reinhardtii* Alb3. The conserved Oxa1/YidC/Alb3 core is shown in yellow, embedded in the thylakoid membrane, while the extended, flexible C-terminal region is shown in coral and projects into the stroma. **d**, Distinct translation states resolved from the complete chlororibosome dataset. Subtomogram averages are shown for elongation factor and tRNA-bound states. tRNAs and translation factors are colored and annotated as indicated.

In the flowering plant Arabidopsis, the protein Alb3, belonging to Oxa1/YidC/Alb3 protein family and also involved in mitoribosome membrane tethering^26–28^, has been shown to be the possible factor involved in co-translational protein insertion in the chloroplast^29,30^. The Chlamydomonas homolog (Uniprot: Q8LKI3) has a disordered and extended C-terminal region that is a strong candidate for mediating ribosome binding, consistent with the flexible connector observed here (Fig. 2c). However, at ∼8 Å resolution for the membrane-bound subclass, the map did not permit unambiguous visualization of density corresponding to this putative C-terminal segment bound to the ribosome. To assess whether the chlororibosomes are translationally active, we performed subtomogram classification and averaging focused on the intersubunit space. This analysis resolved multiple tRNA occupancy states *in situ*, including states with distinct tRNA configurations and translation factors (Fig. 2d). Both free and thylakoid-associated chlororibosomes populate these states, indicating that the membrane-bound complexes are actively translating and engaged in co-translational insertion and/or processing of nascent thylakoid proteins.

The discovery of the SSU arm domain and its presence on translationally active chlororibosomes raised the question of its molecular composition. However, at the resolution of our *in situ* average, we could not unambiguously assign specific polypeptides to this domain.

### Complete composition of the Chlamydomonas chlororibosome solved by single particle cryo-EM

To clarify the identity of the arm domain and obtain a complete model of the Chlamydomonas chlororibosome, we proceeded to solve its structure using cryo-EM single particle analysis (SPA). Chlororibosomes were purified from whole cell lysate using sucrose density gradient centrifugation. Two datasets were collected, one of which was treated with chloramphenicol to stall translation. During image processing (Supplementary Fig. 3), we observed that the additional arm domain density was only resolved in a subset of particles (∼24%), most likely because this module was disturbed during purification. Refinement of all particles yielded a 2.30 Å consensus reconstruction, whereas restricting the analysis to the subset showing the arm domain resulted in a 2.52 Å map that was further improved by focused refinements on different regions of the chlororibosome (Fig. 3a, Supplementary Fig. 3 and 4).

**Figure 3:**
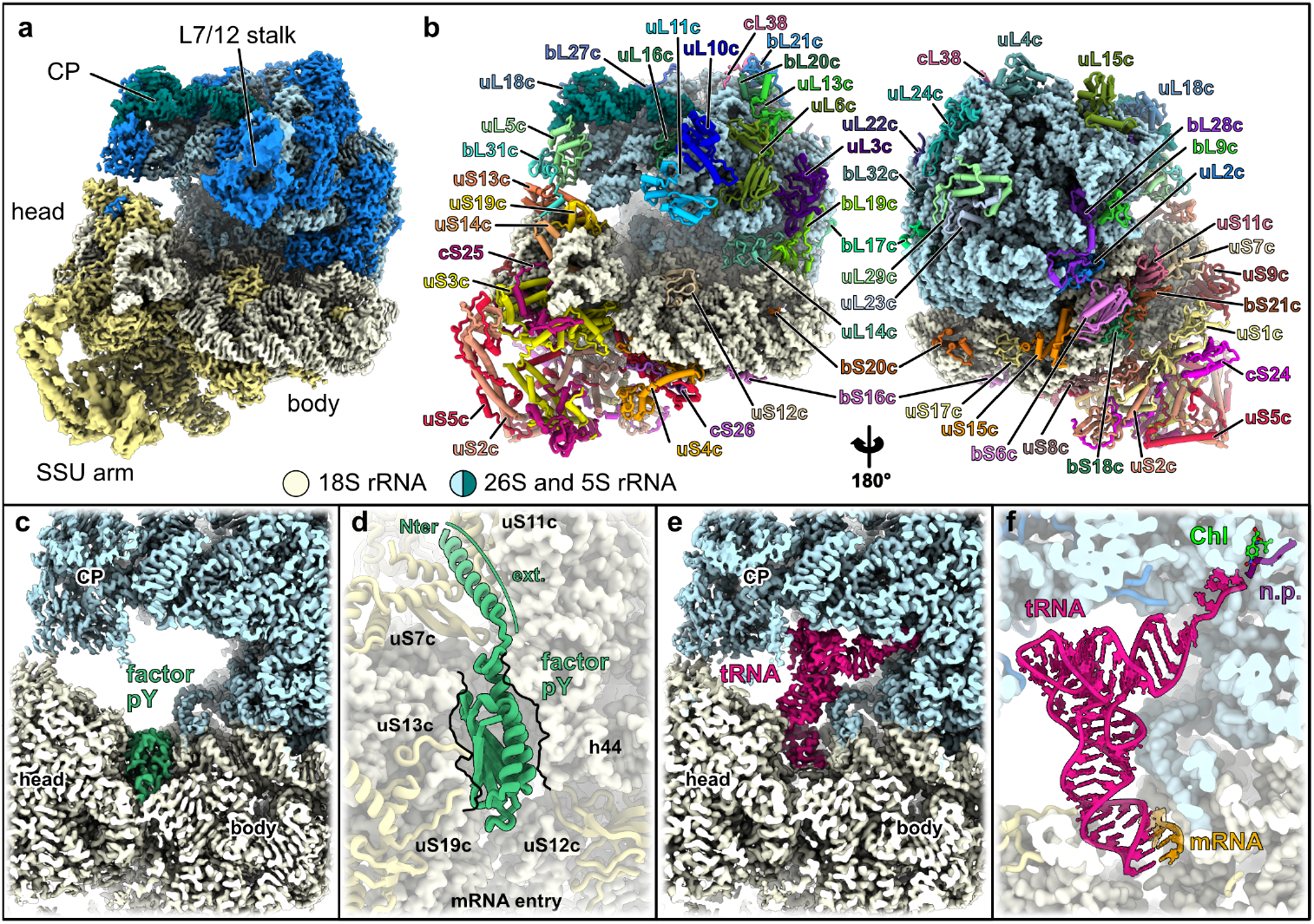
Single particle structure of isolated chlororibosomes. **a**, Consensus composite single particle reconstruction of the *C. reinhardtii* chlororibosome, with components of the SSU shown in beige and components of the LSU shown in blue shades. **b**, Resulting atomic model, with proteins shown in cartoon representation and rRNA shown as surface. All proteins are annotated. **c**,**d**, Zoomed views of the pY-bound chloribosome: density (**c**) and corresponding model (**d**) with pY in green. **d**, pY is shown from the LSU, with the N-terminal extension and the surrounding r-proteins annotated. **e**,**f**, Zoomed views of a bound tRNA: density (**e**) and corresponding model (**f**) with tRNA in magenta, mRNA in orange, and chloramphenicol (Chl) near the nascent polypeptide (n.p.).

This high-resolution SPA map allowed unambiguous identification of a total of 55 proteins, 31 in the LSU and 24 in the SSU, (Fig. 3b) which we consider to be the complete set of core r-proteins of this chlororibosome (see table of proteins, Supplementary Table 2). Compared to a bacterial ribosome such as *E. coli*, the Chlamydomonas chlororibosome possess a complete set of r-proteins except for the absence of bL25 and uL30 from the large subunit (LSU), which was also previously described in Angiosperm chlororibosome structures^14–17^. On the LSU, we find one chloroplast-specific r-protein, cL38/PSPR6, which is conserved with flowering plants, whereas cL37/PSPR5 is not present (Supplementary Fig. 5).

In flowering plants, the chloroplast LSU rRNA is composed of the 5S, 4.5S and 23S rRNAs; the latter is further fragmented post-transcriptionally into three pieces termed A, B and C^14^. In Chlamydomonas, the rRNAs are differentially split (Supplementary Fig. 6). There’s a total of four rRNA pieces: the 5S, 7S, 3S and 23S, which are encoded as such in the chloroplast genome^9^. The 7S rRNA forms most of Domain I, except for helices H19 and H20, which are formed by the 3S rRNA. Helix H18, which would normally connect Domain I to helices H19 and H20, is absent. This leaves the small 3S rRNA fragment stabilized only by r-proteins uL4c and uL24c (Supplementary Fig. 6b). This arrangement helps preserve a bacterial-like architecture in the region surrounding the peptide exit channel. By contrast, the SSU rRNA does not show notable changes from the bacterial organization.

Our cryo-EM map resolution enabled us to visualize rRNA modifications, which are hallmarks of subunit maturation towards a translationally competent ribosome and were not resolved in previous chlororibosome structures. In total, we identified 19 rRNA modifications, 8 in the SSU and 11 in the LSU (Supplementary Fig. 7), all at positions conserved with bacteria. Additional modifications, such as pseudouridine, may be present but could not be resolved, as isomerization modifications generally require ∼2.0 Å resolution for unambiguous identification^31^. Still, this agrees with the fact that the core of the chlororibosome is fundamentally bacterial.

During the analysis, we found that the majority of chlororibosomes were bound to the translation factor pY (also known as PSRP1) (Fig. 3c). This factor was previously described bound to the flowering plant chlororibosome and is involved in light- and temperature-dependent control of protein synthesis^14–17,32–35^. As described in these studies, we observe pY bound in the mRNA channel of the small subunit, where it contacts the 16S rRNA and blocks tRNA binding at both the A and P sites, thereby inhibiting translation (Fig. 3d). pY is present in all previously published chlororibosome structures^14–17^, most likely because it associates with chlororibosomes early during purification, when centrifugation is performed in the dark and subsequent steps at 4 °C. Compared with land-plant pY, the Chlamydomonas factor has a similar core that binds the mRNA channel in an equivalent manner, but it harbors an additional C-terminal extension that projects into the mRNA exit channel (Fig. 3d). To reduce pY binding and obtain a structure of the chlororibosome stalled in a translating state, we also purified them from cells pre-incubated in the presence of chloramphenicol^36^. Thereby, we obtained a stalled chlororibosome structure at 2.84 Å resolution with a P-site tRNA, a segment of mRNA clearly resolved in the mRNA channel, bound chloramphenicol, and the nascent polypeptide chain (Fig. 3e-f). While the mode of interaction of the tRNA, mRNA, and chloramphenicol with the chlororibosome is identical compared to bacteria, this structure provides a stalled translating chlororibosome that we use to investigate the role of the SSU extension (see below).

### The arm domain is primarily formed by large extensions in conserved SSU r-proteins

The most pronounced differences at the protein and structural levels are found in the SSU. Relative to bacterial and plant chloroplast ribosomes, the SSU contains a nearly complete set of canonical r-proteins, with the notable exception of bTHX, which is absent here while present in the Chlamydomonas mitochondrial ribosome^26^. In addition, chloroplast-specific r-proteins cS22 and cS23 (PSRP2 and PSRP3), which in flowering plants localize to the SSU foot^14^, are entirely missing (Supplementary Fig. 5). Together, these features render the SSU strongly bacterial-like, apart from the extended SSU arm domain that projects from this conserved core (Fig. 4).

**Figure 4:**
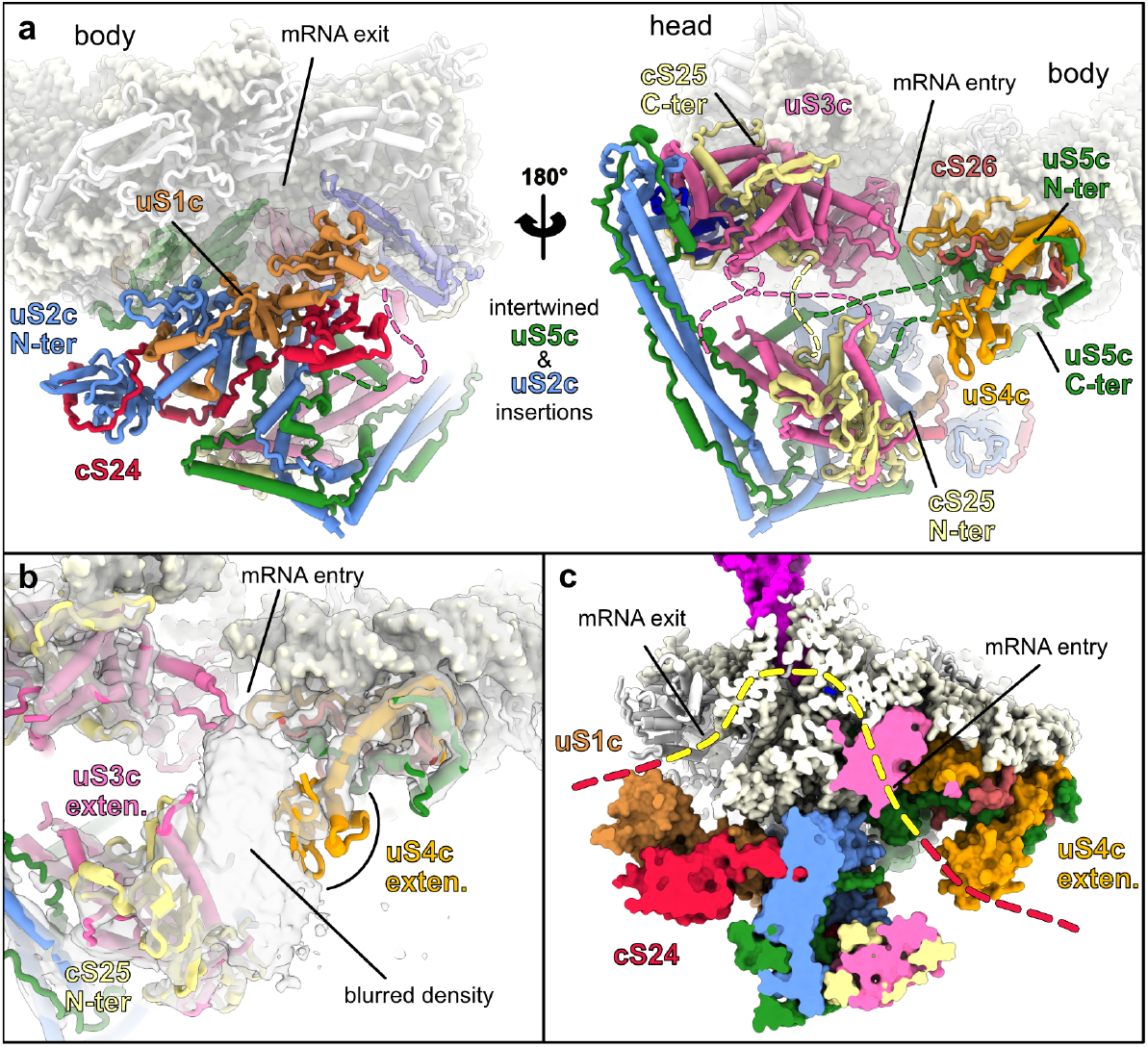
The SSU arm domain extends the entry and exit of the mRNA channel. **a**, Overall architecture of the SSU arm. Two views related by a 180° rotation show the arrangement of arm components from the mRNA exit and entry. Proteins are shown in cartoon representation. **b**, Close-up of the mRNA entry site. The model is overlayed with the map of the tRNA class. A blurred density is sandwiched between the uS4c extension and the cS25 N-terminus, in proximity to the uS3c extension, consistent with an averaged segment of mRNA. **c**, Cut view of the SSU highlighting the prolonged mRNA path shaped by the arm domain. The canonical path is indicated in yellow (dashed), whereas the prolonged path observed here is shown in red (dashed). The mRNA entry is stabilized mainly by the uS4c extension, and the exit by uS1c, which itself is stabilized by cS24.

The SSU arm is positioned on the solventexposed side of the small subunit, anchored around the mRNA entry and exit channels (SSU back) of the SSU body, from which a long, straight segment extends toward the SSU head. It is formed by extensions and insertions in the r-proteins uS2c, uS3c uS4c and uS5c, which are all encoded in the chloroplast genome, together with three chloroplast-specific r-proteins, PSRP3, PSRP7, and the previously uncharacterized PSRP8, which are not present in flowering-plant chlororibosomes. Following ribosomal protein nomenclature^37^, and because they are core components of the Chlamydomonas chlororibosome, we refer to these additional proteins as cS24, cS25 and cS26, respectively.

cS24/PSRP3 is located at the mRNA exit channel on the back of the SSU, where it stabilizes the interface between the conserved cores of bS1c and uS2c (Supplementary Fig. 5). cS25/PSRP7 is the largest (560 aa) of the chlororibosome-specific proteins. Its C-ter part is positioned on the beak region of the SSU head, where it is intertwined with the extensions of uS3c and a portion of uS10c, while its N-ter part extends to the core of the SSU arm domain. On the body, cS26/PSRP8 functionally replaces the C-ter portion of uS4c, which in Chlamydomonas has an extension that forms a specific domain. There, cS26/PSRP8 interacts with the core parts of uS4c and uS5c. Together, these chloroplast-specific proteins appear to stabilize regions of the chlororibosome that harbor extensive r-protein extensions.

Most of the SSU arm domain, however, is contributed by extensions in uS2c, uS3c, uS4c and uS5c, which was previously hinted by early proteomic studies^38–40^. Relative to their bacterial counterparts, uS2c, uS3c, uS4c and uS5c are extended by 666, 497, 97 and 512 residues, respectively (Supplementary Fig. 8). The relatively small C-terminal extension of uS4c folds into a globular domain near the mRNA entry channel. By contrast, uS2c is the protein with the largest extension. On its N-ter, the extension forms three S1-like β-barrel domains that run along the SSU body in proximity to uS8c. Only the third (most C-terminal) S1-like domain could be fully modeled, stabilized by interactions with cS24/PSRP3 (Supplementary Fig. 9). The first two S1-like domains are poorly resolved in SPA reconstructions but are clearly visible in the native STA map (Supplementary Fig. 9c). The insertion of uS2c (440-795 aa) consists mainly of alpha helices that form a coiledcoil extending away from the SSU back, where it intertwines by charge complementarity with the unstructured extension of uS5c (Supplementary Fig. 8a,c). The intertwined uS2c/uS5c arm reaches toward the SSU head to contact part of the extension made by uS3c and is further stabilized by the N-terminal segment of uS10c. Because only a few residues participate in this interface, the interaction is likely weak, and the entire uS2c/uS5c arm may readily unlatch. This would explain why the native conformation was retained in only a subset of chlororibosomes visualized by SPA; the arm domain likely unlatched during isolation and was therefore too flexible to be resolved in the average. Finally, uS3c extensions enlarge the SSU beak region but also projects at the base of the intertwined uS2c/uS5c segment. There, it is made of several alpha helices that interweave with the N-ter part of cS25/PSRP7 to form a platform-like structure (Fig. 4). Using Foldseek^41^, we analyzed the conservation of the SSU arm. Analysis worked best on uS2c and uS3c, which have the most defined folds, and shows that this domain is restricted to Chlorophyceae (Supplementary Fig. 10).

Looking at the SPA reconstructions of chlororibosomes bound to the P-site tRNA, we observed a blurred density near the SSU arm domain, positioned between the uS4c extension and the intertwined cS25 and uS3c extensions (Fig. 4b). This density most likely corresponds to averaged mRNAs stalled on the ribosome, which are expected in the tRNA class. The uS4c and uS3c extensions positioned there are overall positively charged (Supplementary Fig. 8b), which is compatible with a role in mRNA stabilization. These observations suggest that the uS4c extension and the platform made of cS25 and uS3c extensions help stabilize and/or thread the mRNA at the mRNA entry channel. On the other side, at the mRNA exit, the protrusion formed by cS24/PSRP3, uS2c, and bS1c (Fig. 4a) also points towards stabilization of the mRNA at the exit channel (Fig. 4 and Supplementary Fig. 9). Together, the rprotein extensions and small proteins of the arm domain appear to prolong the mRNA path, at both the mRNA entry side (uS4c, uS3c, cS25) and the mRNA exit side (cS24/PSRP3, uS2c, bS1c) of the chlororibosome SSU (Fig. 4c).

In summary, our SPA and *in situ* cryo-ET reveal that Chlamydomonas chlororibosomes adopt a dynamic mode of membrane engagement and harbor an accessory arm domain on the SSU, an unexpected discovery that refutes the previous assumption that all chlororibosomes are structurally similar. In Chlamydomonas, membrane-bound chlororibosomes are only loosely tethered to thylakoids via a thin connector that is consistent with a flexible Alb3–nascent chain interaction, allowing substantial orientational freedom rather than rigid docking at the translocon. The SSU arm domain emerges as a noticeable architectural innovation: it is built predominantly from chloroplast-encoded extensions and insertions in canonical SSU r-proteins, together with three chloroplast-specific r-proteins. Its position decorates the mRNA entry and exit channels (Fig. 4c) suggesting roles in stabilizing the mRNA and organizing chloroplast polysomes. Additionally, the long arm structure projecting from the back toward the head might play a role in recruiting regulatory factors, although the precise binding partners remain unknown. Supporting this hypothesis, several gene-specific *trans*-acting proteins have been described to be involved in mRNA stabilization and required for efficient translation in the Chlamydomonas chloroplast^42–44^. Given that Chlamydomonas is relatively close to land plants on the green lineage, our findings imply that chlororibosomes are likely more structurally and functionally diverse than previously appreciated, with potentially even greater elaboration in secondary plastid-bearing algae that dominate marine phytoplankton and remain to be explored.

## Acknowledgements

We thank M. Chami and D. Kalbermatter from the University of Basel BioEM facility for their help operating the electron microscopes during the cryo-EM sample screening. We thank F. Wilmund, R. Bock, D. Sloan and W. Wietrzynski for the scientific discussions. We thank AG. Seviné and M. Zavolan for providing access to the BioComp gradient fractionator.

## Author contributions

F.W. and B.D.E conceptualized this study. S.P. performed initial subtomogram averaging. F.W. and P.A.L. carried out ribosome purification and single-particle cryo-EM data processing. F.W. performed cryo-electron tomography data processing, model building, and data interpretation. Singleparticle cryo-EM data collection was performed by L.K. Membrane movement analyses were conducted by P.V.d.S. The manuscript was written by F.W. and B.D.E., with input from all authors.

## Data and material availability

The single particle cryo-EM maps of C. reinhardtii chlororibosome have been deposited at the Electron Microscopy Data Bank (EMDB) and models on the protein data bank (PDB). For the overall single particle reconstruction, composite map EMD-56352 (PDB: 9TVU), consensus map EMD-56337, LSU focus EMD-56338, LSU CP focus EMD-56339, LSU L7/12 focus EMD-56340, SSU body EMD-56341 and SSU head EMD-56342, SSU extension base EMD-56349 and SSU extension tip EMD-56344.

For the SPA in presence of the factor pY composite map EMD-56602 (PDB: 28LU), consensus map EMD-56560, LSU focus EMD-56563, SSU focus EMD-56564. For the SPA with the P-site tRNA composite map EMD-56567 (PDB: 28JW), consensus map EMD-56561, LSU focus EMD-56562, SSU focus EMD-56565.

For the subtomogram average of the native Chlamydomonas chlororibosome: composite map EMD-56143, consensus map EMD-56139, LSU focus EMD-56140, SSU focus EMD-56141 and SSU extension EMD-56142, and membrane-bound class EMD-56144. The raw cryo-ET data is available at the Electron Microscopy Public Image Archive (EMPIAR), accession code: EMPIAR-11830.

## Materials and Methods

### Cryo-ET data acquisition and preprocessing

All data analysis was performed on the cryo-ET dataset deposited under EM-PIAR-11830^23^. Briefly, tilt-series data were collected using a Titan Krios G4 transmission electron microscope operating at 300 kV equipped with a Selectris X energy filter with a slit set to 10 eV and a Falcon 4i direct electron detector (Thermo Fisher Scientific) recording dose-fractionated movies in EER format. A dose-symmetric tilt scheme using TEM Tomography 5 software (Thermo Fisher Scientific) was employed in the acquisition with a tilt span of ± 60°, covered by 2° or 3° steps starting at a 10º pre-tilt. Target focus was set for each tilt-series in a range of -1.5 µm to -3.5 µm in steps of 0.25 µm. The microscope was set to a magnification corresponding to a pixel size of 1.96 Å at the specimen level and a nominal dose of 3.5 e^-^/Å^2^ per tilt image.

### Subtomogram averaging

129 chloroplast tomograms (Supplementary Table 3) from the Chlamydomonas dataset^23^ were selected based on overall quality and the abundance of chloroplast ribosomes. Data processing followed the workflow described in the TomoGuide^44^ using a post-acquisition refined pixel size of 1.91 Å^23,45^. See Supplementary Fig. 1 for processing workflow. Using AreTomo3^46^, raw EER movies were automatically motion-corrected, their CTF estimated, tilt series aligned, and CTF-corrected tomograms reconstructed. Denoising was performed with Icecream on tomogram pairs reconstructed from odd or even raw frames respectively^47^.

For particle picking, 3D template matching was carried out on bin4 CTF-corrected tomograms (7.64 Å/pixel) with 6° angular sampling in pyTOM-match-pick^48^, using a chloroplast ribosome reference obtained from single-particle analysis and low-pass filtered to bin4 Nyquist (15.28 Å). For each tomogram, the number of extracted positions were automatically determined by pytom and the threshold was reduced by 5% to maximize the number of true positives, yielding an initial set of 20,883 picks. Picks were then imported in RELION5^49^ and the particles were cleaned by several rounds of 3D classification without alignment using bin4 subtomograms, to clean out obvious false positives (membranes and high signal contaminants). At this step, the 14,415 remaining particles were 3D refined and reached bin4 Nyquist (15.28 Å).

From there, a bin1 reference (8.87 Å) was generated for CTF refinement and Bayesian polishing, improving the overall resolution to 6.93 Å. Particles were reextracted at bin2, 3D refined to 7.64 Å, and cleaned by an additional round of 3D classification, resulting in 13,245 particles. A new bin1 reference (6.79 Å) was used for a second round of CTF refinement and Bayesian polishing, yielding a 6.70 Å map. Finally, particles were reextracted at bin1, refined to 6.39 Å, followed by a final round of CTF refinement and Bayesian polishing, which resulted in a 6.11 Å reconstruction. Focused refinements using masks on the LSU, the SSU, and the SSU-extension yielded maps at 5.76 Å, 6.36 Å, and 8.48 Å, respectively.

Additionally, particles were reextracted at bin3 to classify for tRNA states and membrane-bound ribosomes. For membrane-bound ribosomes, 3D classification was performed without alignment in two successive rounds using a mask specifically covering the membrane area of the particles. This yielded a subset of 4,160 particles, which were used to generate the final 9.29 Å resolution reconstruction. For the tRNA states analysis, a similar approach was used using a mask covering the tRNA region. Particles were 2 times classified using a high T parameter (T=10) which resulted in 6 distinct states.

### Membrane movement analysis

From the membrane-bound ribosomes, particles were further 3D classified without alignment using a masks pecifically covering the membrane area. Using two different T parameters (T= 2 or 4), 8 different membrane orientation classes were obtained. From there orientations of the ribosomes relative to their membranes were quantified by finding the 3D normal vector of the plane fitted to the membrane density. Percentile thresholding of the voxel intensity in an annular region was used for membrane identification. The normal vector was determined through principal components of the 2D in-plane orientation and interpolated by linear regression across the Z-slices. Angular deviations from the mean normal orientation were projected onto a 2D tangent plane and visualized as a kernel density map. Full script available at https://github.com/Phaips/Ribo-Move.

### Purification of choloroplastic ribosomes

Chloroplast ribosomes were purified from 2L of Chlamydomonas culture (WT2.2 strain), grown under continuous light in TAP media. Cells were harvested by centrifugation at 2000g for 5 min at room temperature. They were then resuspended in 30mL Lysis buffer (20mM HEPES-KOH – pH7.6, 75mM KCl, 15mM MgCl2, 1.5mM DTT, C0mplete EDTA-Free) and cells were ruptured using a French Press by 2 passes at XX pressure (900). 1% (final concentration) of Triton-X100 was added to the lyzed cells. The solution was transferred to a 40 mL glass Dounce potter and homogenized by 10 strokes to ensure complete disruption of the chloroplast and solubilization of the membranes. Cell debris were removed by two rounds of centrifugation at 500g for 5min. From there, ribosome purification followed the same procedure as previously described^50^. The supernatant was loaded onto a 40% sucrose cushion in monosome buffer (same as lysis buffer, but without Triton X-100) and centrifuged at 235,000g for 3 h at 4 °C. The crude ribosome pellet was resuspended in monosome buffer and loaded onto a 10–30% sucrose gradient prepared using the BioComp gradient master 108, and run for 16 h at 65,000g in SW41 rotor. Fractions corresponding to chlororibosomes were collected using a BioComp piston gradient fractionator, pelleted and resuspended in monosome buffer. For the chloramphenicol stalled ribosomes, cells were incubated with chloramphenicol for 1 hour at a concentration of 500 µg/mL before cell lysis, and 100 µg/mL was kept throughout the entire purification processes.

### Grid preparation

4 µl of chloroplast ribosomes at a concentration of 3 µg/µl of proteins (BSA equivalent measurement on Nanodrop) were applied onto a Quantifoil R2/1 300-mesh holey carbon grid coated with 2nm continuous carbon film that were glow-discharged at 5W for 20 sec in a Gatan glow discharger. The sample was plunge frozen in liquid ethane with a Vitrobot Mark IV system (temperature = 4 °C, humidity 100%, blot force 5) with a wait time of 25 sec before blotting for 2.5 sec.

### Single particle data collection

Data collection was performed using a 200kV Glacios and a 300kV Titan Krios electron microscope (Thermo Fisher Scientific) both equipped with Falcon4 camera using EPU (Thermo Fisher Scientific) for automated data acquisition. Data were collected at a nominal underfocus of −0.8 to −2.0 µm at a magnification of ×120k for the Glacios and ×75k for the Krios, both yielding a pixel size of 0.84 Å. For the 200kV dataset, 15,628 micrographs were recorded as a.tif movie stack, each movie stack was fractionated into 50 frames for a total exposure corresponding to an electron dose of 49 e^-^/Å^2^. For the 300kV dataset, 20,779 micrographs were recorded as a.tif movie stack, each movie stack was fractionated into 51 frames for a total exposure corresponding to an electron dose of 50 e^-^/Å^2^.

### Single particle cryo-EM data processing

For both datasets, all single-particle processing was carried out in cryoSPARC^51^, and the workflows are summarized in Supplementary Fig. 3 and 4. The 200 kV and 300 kV datasets were pre-processed, particles picked, classified, and refined independently; only at the final stage were selected particles from the two datasets merged. After motion correction and CTF estimation, 15,491 micrographs (Dataset1: 200 kV) and 19,406 micrographs (Dataset2: 300 kV) were retained.

For the 200 kV dataset, particles were picked using Template Picker, extracted in a 648-pixel box downsampled to 256 pixels (2.126 Å/pixel), and cleaned by two rounds of 2D classification, yielding 936,493 particles. Subsequent 3D cleaning using Heterogeneous Refinement and 3D classification produced 491,158 good particles. For the 300 kV dataset, an equivalent pipeline yielded 456,710 good particles after 3D classification. Chlororibosomes were then classified based on the presence of the SSU extension using a mask encompassing the entire SSU and the extension, using a combination of 3D classification and 3D variability analysis. This yielded 129,415 particles with the SSU extension in the 200 kV dataset and 97,236 particles with the SSU extension in the 300 kV dataset.

Particles were re-extracted for high-resolution refinement in a 648-pixel box downsampled to 512 pixels (1.063 Å/pixel). For the 200 kV dataset, the SSU extension class reached 3.10 Å after global refinement, improved to 2.76 Å after combined global and local CTF refinement, and finally to 2.60 Å after reference-based motion correction followed by another round of global and local CTF refinement. For the 300 kV dataset, the SSU extension class reached 3.40 Å after global refinement, improved to 2.88 Å after global and local CTF refinement, and 2.71 Å after reference-based motion correction plus global and local CTF refinement.

At this stage, particles from the SSU extension classes of both datasets were merged, resulting in 225,199 particles that refined to 2.52 Å resolution. Focused refinements were then performed using soft masks on the LSU (2.45 Å), followed by local refinements of the LSU L7/12 stalk (3.99 Å) and central protuberance (2.55 Å). Analogous focused refinements were carried out for the SSU (2.63 Å), including separate refinements of the SSU head (2.63 Å), SSU body (2.57 Å), and the SSU extension base (2.98 Å) and tip (3.94 Å).

From these, 175,058 particles had a clear pY factor and 49,141 a P-site tRNA. The P-site tRNA class was refined to 2.84 Å resolution and was further focused refined on the LSU (2.76 Å) and the SSU (2.88 Å). The pY factor class was refined to 2.56 Å resolution and was further focused refined on the LSU (2.48 Å) and the SSU (2.63 Å).

### Model building and refinement

Focused refined cryo-em maps (all at resolution better than 3 Å) for the large subunit, body and head of the small subunit were used as inputs for ModelAngelo^52^ *de novo* sequencing from the experimental data (without input sequence). Peptide chains obtained were then blasted against *Chlamydomonas reinhardtii* UNIPROT database (Taxon ID 3055) and the corresponding AlphaFold2^53^ models were retrieved for all the identified proteins (all were identified). AlphaFold2 models were then matched to the ModelAngelo models using the Matchmaker tool in ChimeraX^54^, and then further rigid-body fitted in their respective cryo-em maps. The protein models were then manually inspected and refined in COOT^55^. For the rRNAs, the models from *E. coli* 8B0X^56^ were rigidbody fitted in the density and then refine with restraints in COOT. The sequence was then mutated accordingly and refined again. For ions, ligands and rRNA modification, they were first placed by homology with 8B0X and then manually curated in COOT. The different parts of the model were then automatically refined in PHENIX against the best resolved focus-refined maps using the phenix.real_space_refine tool and then again manually refined in COOT, this repeating through several cycles. The model geometry was validated using MolProbity^57^ and is summarized in Supplementary Table 1.

### Figure preparation

All figures were prepared using UCSF ChimeraX, developed by the Resource for Biocomputing, Visualization, and Informatics at the University of California, San Francisco, with support from National Institutes of Health R01-GM129325 and the Office of Cyber Infrastructure and Computational Biology, National Institute of Allergy and Infectious Diseases^58,59^. For tomogram views with map-backs particle map back, the ArtiaX plugging was used^60^ and membrane segmentations for visualization were automatically performed using MemBrain-seg^61^.

