## Supplementary Fig. 1 for "Chloroplast-encoded small subunit extensions reshape the Chlamydomonas chlororibosome"

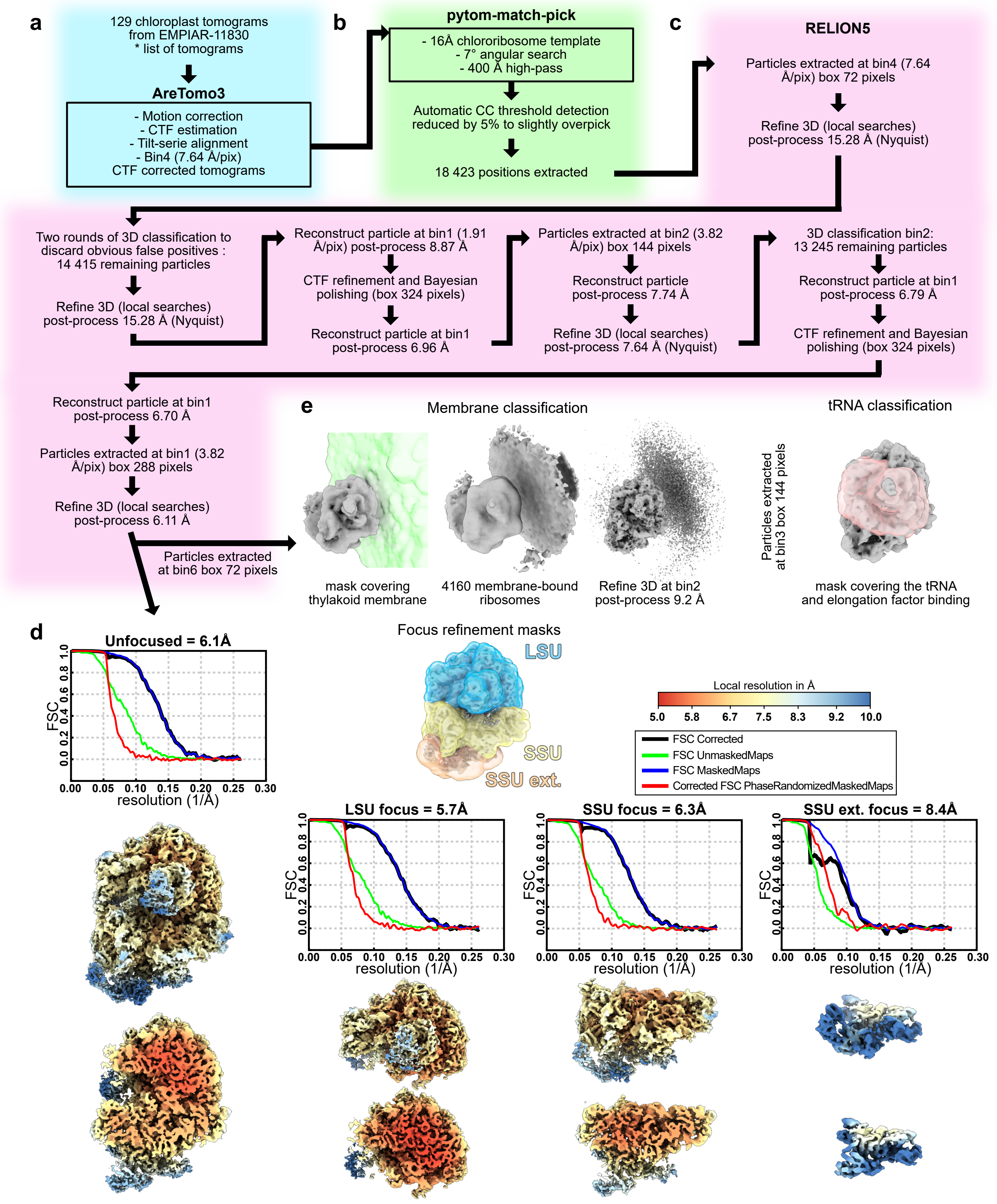


**Supplementary Figure 1:** Tomography data processing summary

**a** Tilt-series processing and tomogram reconstruction from 129 chloroplast tomograms (EMPIAR-11830, see Supplementary Table 3) using AreTomo3, including motion correction, CTF estimation, tilt-series alignment, and reconstruction of CTF-corrected tomograms (shown at bin4; 7.64 Å/pixel). **b** Particle picking by 3D template matching in pyTOM-match-pick on bin4 tomograms using a chloroplast ribosome SPA reference low-pass filtered to bin4 Nyquist (15.28 Å), with 6° angular sampling and a 400 Å high-pass. **c** Particle extraction, 3D classification/cleaning, and iterative refinements in RELION5, including re-extraction at higher sampling and CTF refinement/Bayesian polishing. Focused refinements of the LSU, SSU, and SSU-extension were performed using the masks shown. Gold-standard FSC curves are plotted with resolution reported at the FSC = 0.143 criterion. Local-resolution estimates are displayed on the corresponding maps using a common color scale; the local-resolution range for each map is indicated below. Additional 3D classifications used to separate tRNA states and membrane-bound ribosomes (mask-restricted, without alignment) are part of the processing but are not depicted in this schematic.
