## Supplementary Fig. 2 for "Chloroplast-encoded small subunit extensions reshape the Chlamydomonas chlororibosome"

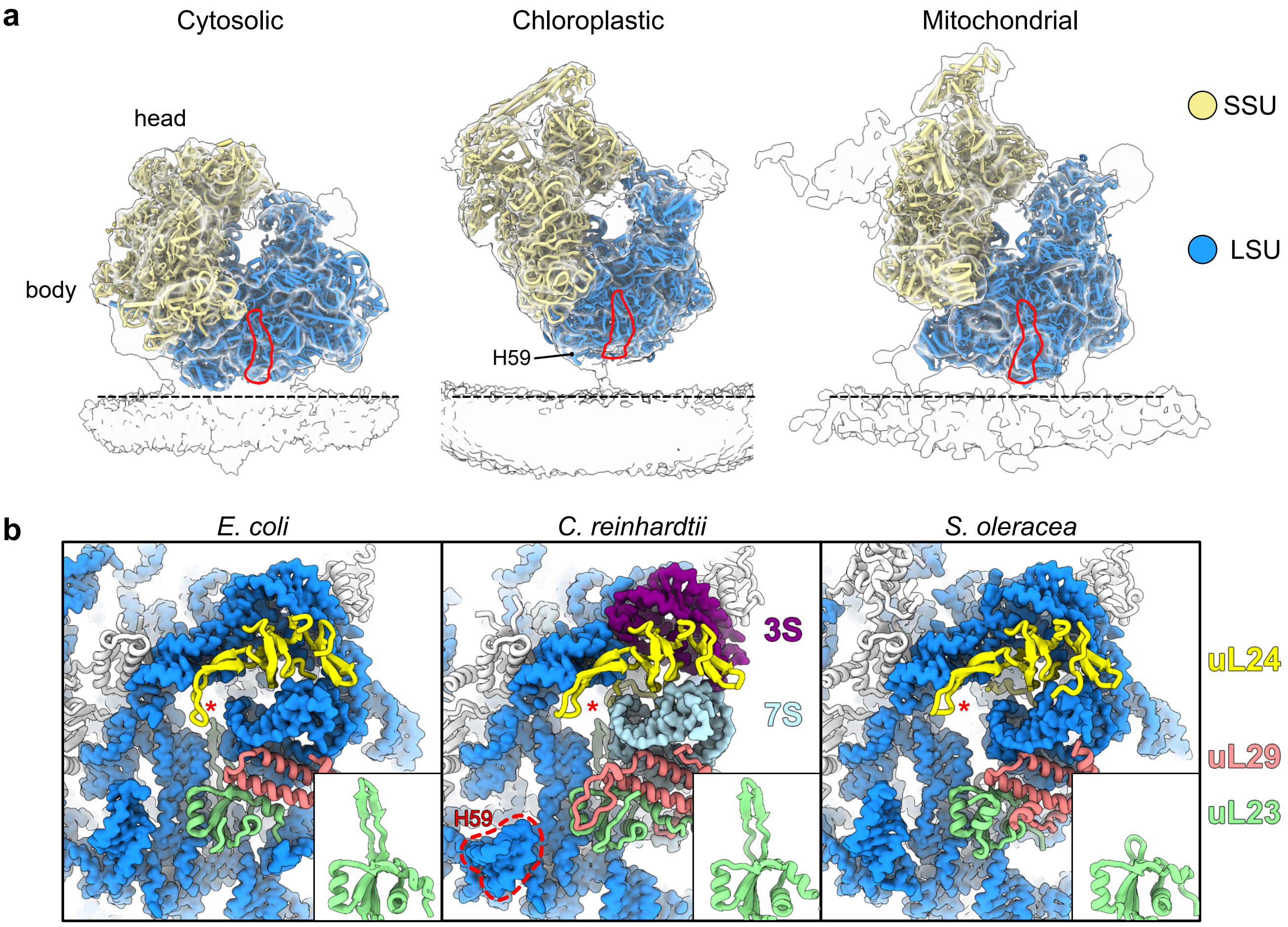


**Supplementary Figure 2:** Comparative analysis of the membrane association

**a** Comparison of the membrane association between the cytosolic, chloroplastic and mitochondrial ribosomes in Chlamydomonas. The peptide channel path is highlighted in red. **b** Comparison of the Chlamydomonas chlororibosome area surrounding the peptide exit channel with the E. coli ribosome (PDB: 8B0X) and Spinach chlororibosome (PDB: 6ERI). rRNAs are shown in surface representation and r-proteins in cartoon. The insert shows uL23, notably the shortened loop in the Spinach chlororibosome, that is not occurring in Chlamydomonas.
