## Supplementary Fig. 3 for "Chloroplast-encoded small subunit extensions reshape the Chlamydomonas chlororibosome"

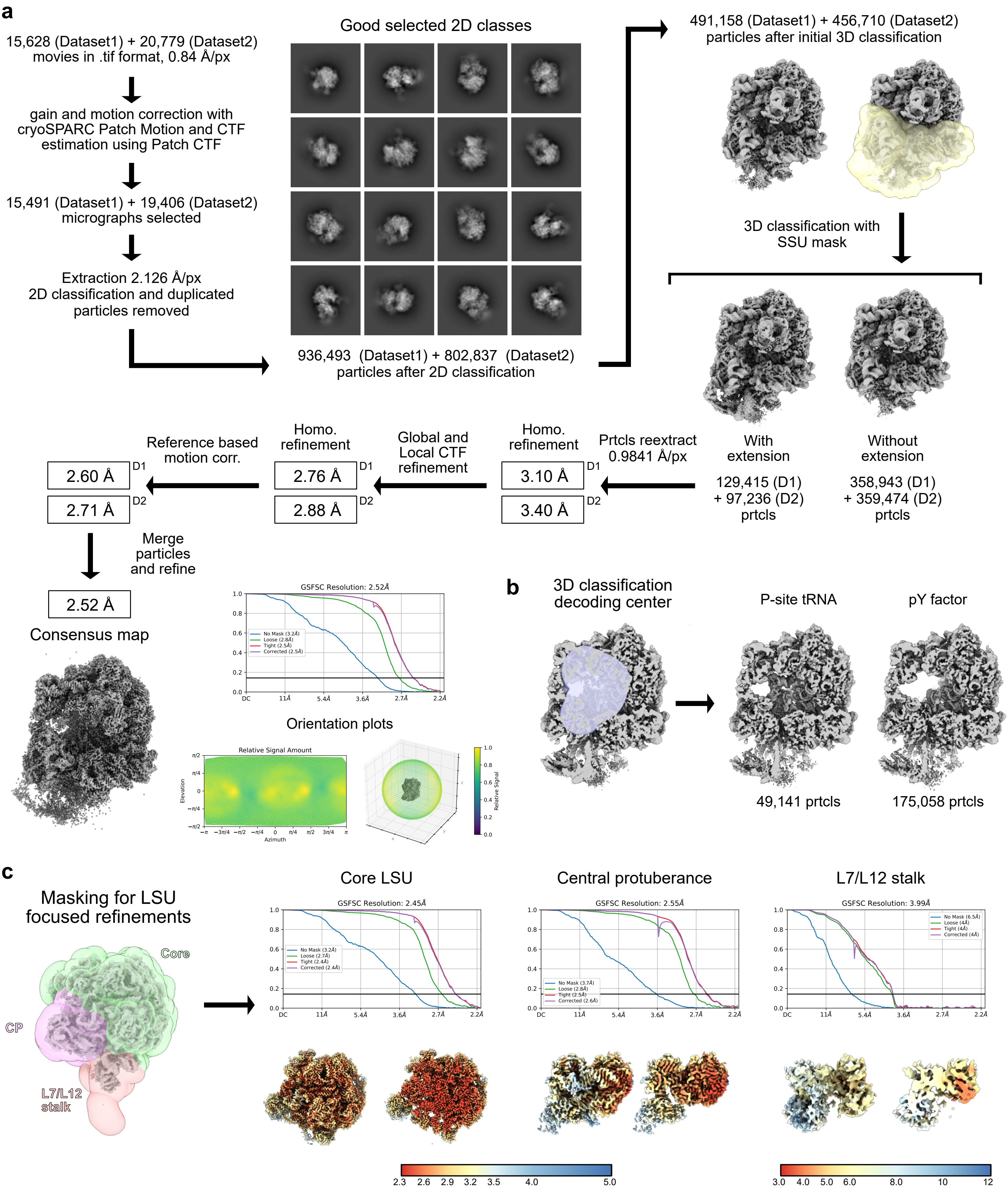


**Supplementary Figure 3:** Single-particle data processing workflow

Graphical summary of the processing workflow described in Methods. **a** Post-processing, 2D and 3D classification. An orientation plot generated from the Orientation Diagnostic cryoSPARC tool is shown. **b** 3D classification for the decoding area. **c** Focused refinements for the LSU, with all the masks used shown. For the final reconstruction, GSFSC curves are plotted, with resolution calculated at the 0.143 threshold. Local resolution plotted on the maps were generated using the built-in cryoSPARC tool with default parameters, all using the same resolution scale, with maps also shown in cut view.
