## Supplementary Fig. 4 for "Chloroplast-encoded small subunit extensions reshape the Chlamydomonas chlororibosome"

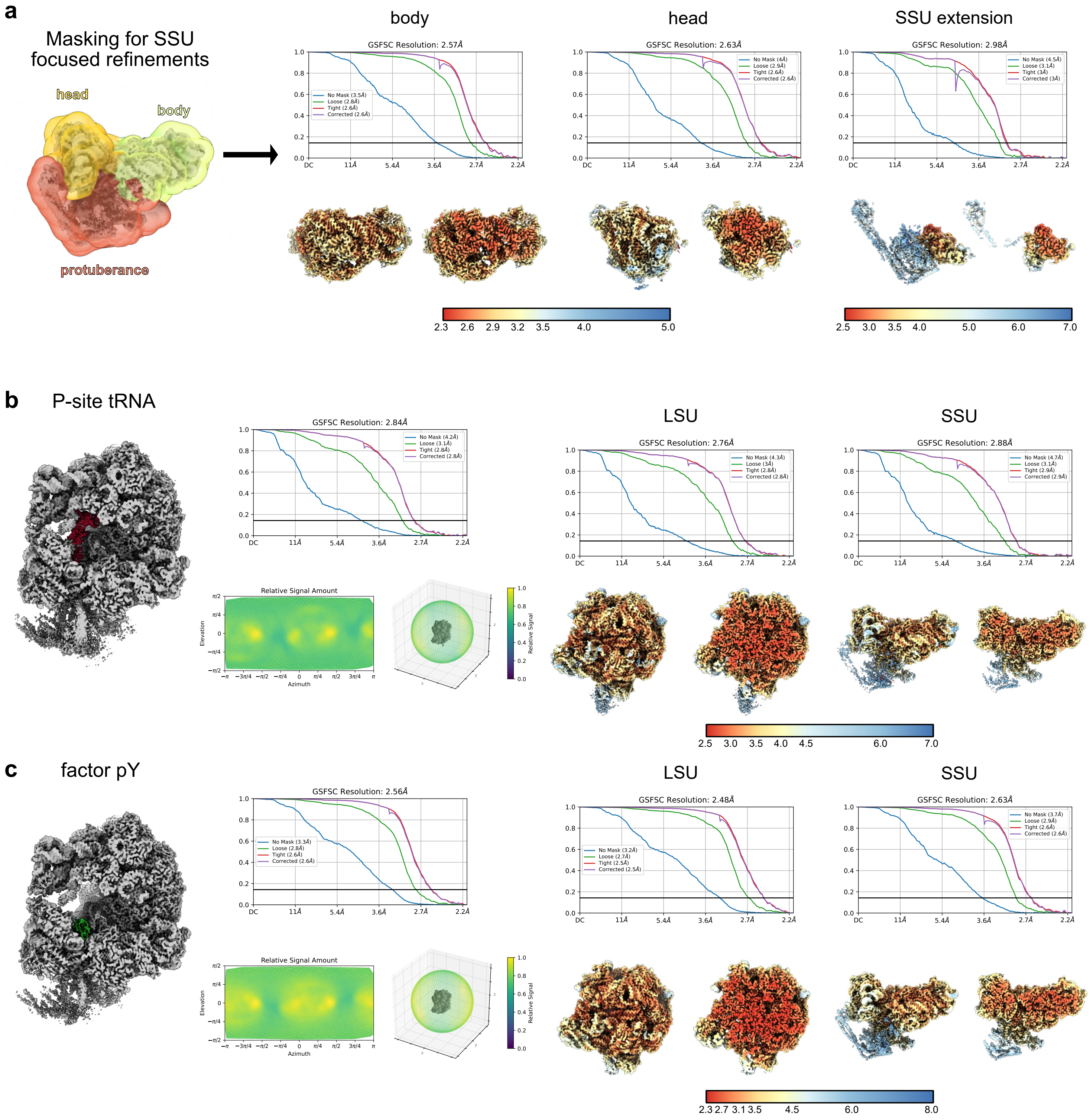


**Supplementary Figure 4:** Single-particle data processing workflow, FSC plots of the SSU and tRNA and pY states

**a** Focused refinements for the SSU of the consensus map, with all the masks used shown. For the final reconstruction, GSFSC curves are plotted, with resolution calculated at the 0.143 threshold. Local resolution plotted on the maps were generated using the built-in cryoSPARC tool with default parameters, with a resolution scale different for the SSU head extension, with maps also shown in cut view. **b** and **c** show the consensus refinements for the P-site tRNA and pY factor classes with cryoSPARC orientation plots shown, as well as the respective focused refinements done on the LSU and SSU.
