## Supplementary Fig. 5 for "Chloroplast-encoded small subunit extensions reshape the Chlamydomonas chlororibosome"

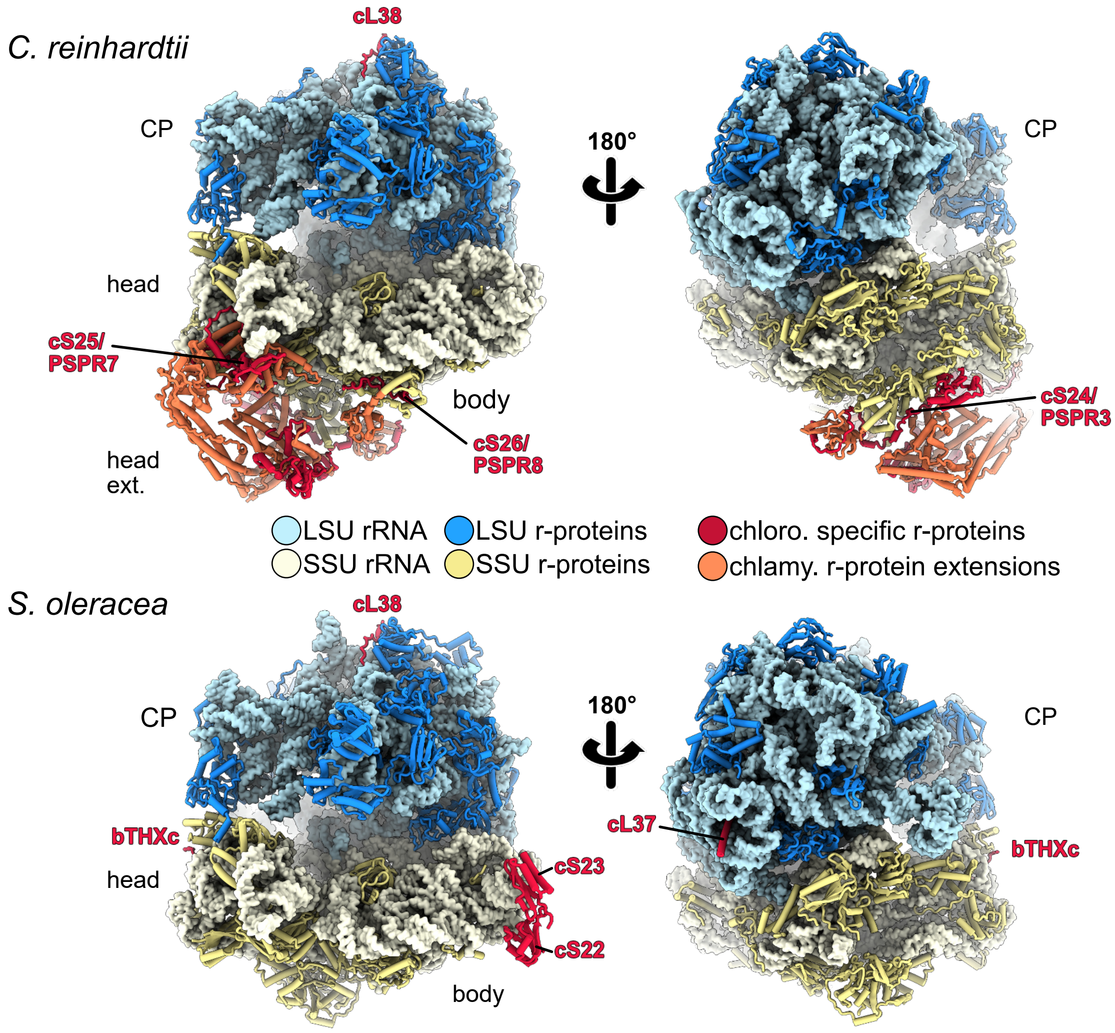


**Supplementary Figure 5:** Comparison specific r-proteins between Chlamydomonas and Spinach

Chlororibosomes from *C. reinhardtii* (top) and *S. oleracea* (bottom; PDB: 5MMM) are shown in two views related by a 180° rotation. rRNA are shown in surface representation and r-proteins in cartoon and color key is indicated on the figure. Chloroplast-specific r-proteins are highlighted in red and *Chlamydomonas*-specific r-protein extensions in coral. Features of the ribosome are labeled, with CP for central protuberance; "head", "body", and "head ext.".
