## Supplementary Fig. 6 for "Chloroplast-encoded small subunit extensions reshape the Chlamydomonas chlororibosome"

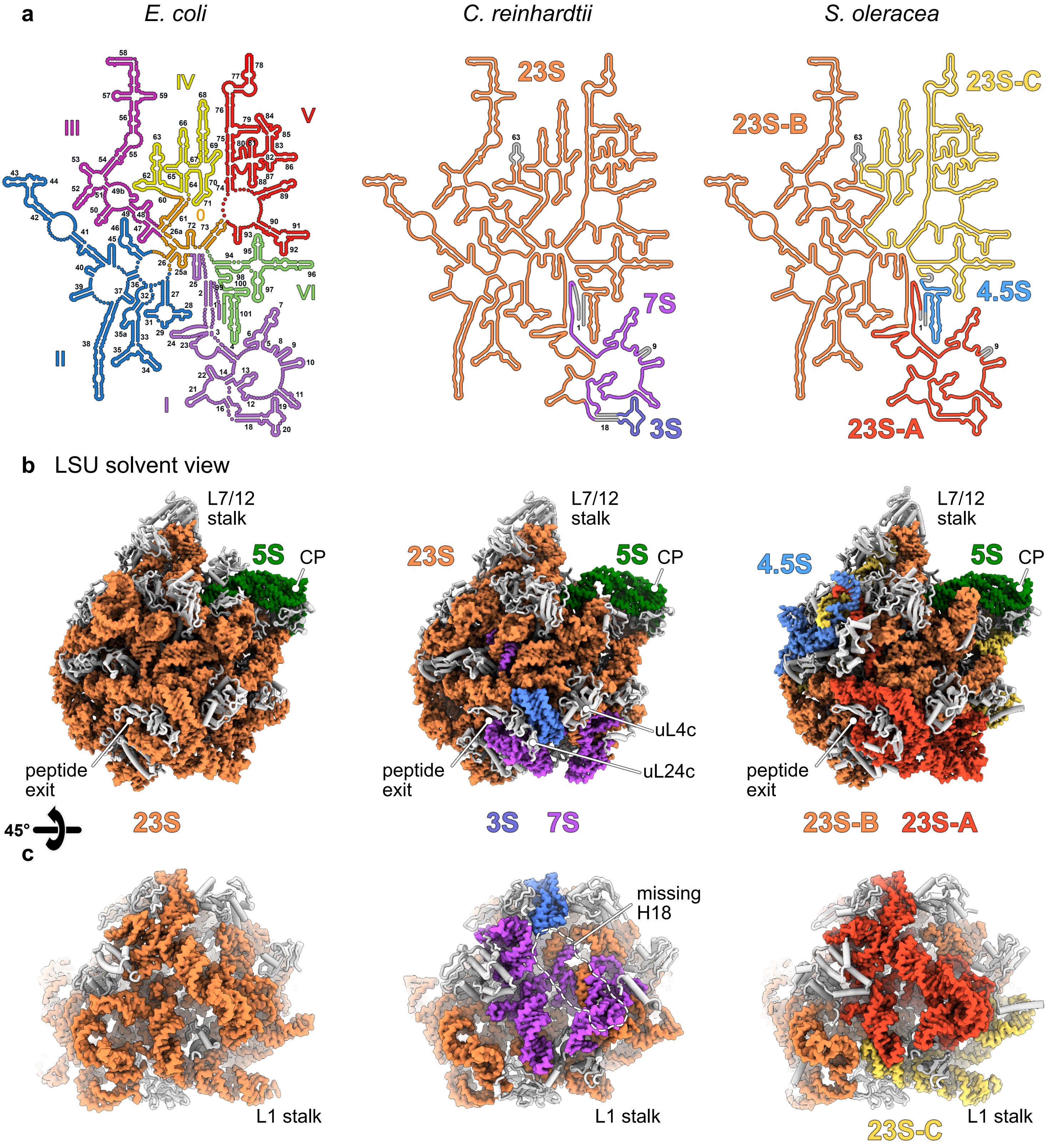


**Supplementary Figure 6:** Fragmentation of the LSU's rRNAs

**a** 2D diagrams of the LSU rRNAs from *E. coli*, *C. reinhardtii* and *S. oleracea*. The E. coli one is colored by domains with all helices annotated, while the *C. reinhardtii* and *S. oleracea* ones are colored according to the rRNA fragments with missing helices annotated and grayed out. **b** and **c** shows two different view of the LSU in 3D with rRNAs colored according to **a** and r-proteins shown in white.
