## Supplementary Fig. 7 for "Chloroplast-encoded small subunit extensions reshape the Chlamydomonas chlororibosome"

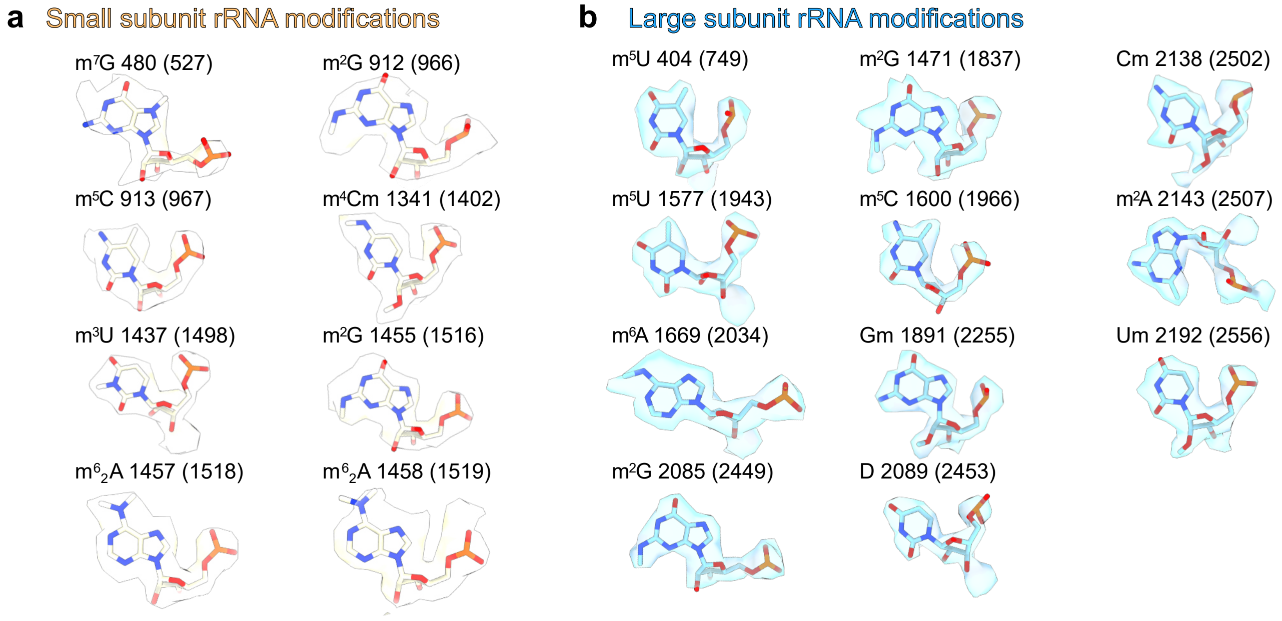


**Supplementary Figure 7:** Chlamydomonas chlororibosome rRNA modifications

All the rRNA modifications identified from the cryo-EM map are shown in their respective densities with modifications in **a** the LSU 26S rRNA shown in light blue and in **b** the SSU 18S rRNA shown in beige. Numbering in *E. coli* (according to PDB: 8B0X) is shown.
