## Supplementary Fig. 8 for "Chloroplast-encoded small subunit extensions reshape the Chlamydomonas chlororibosome"

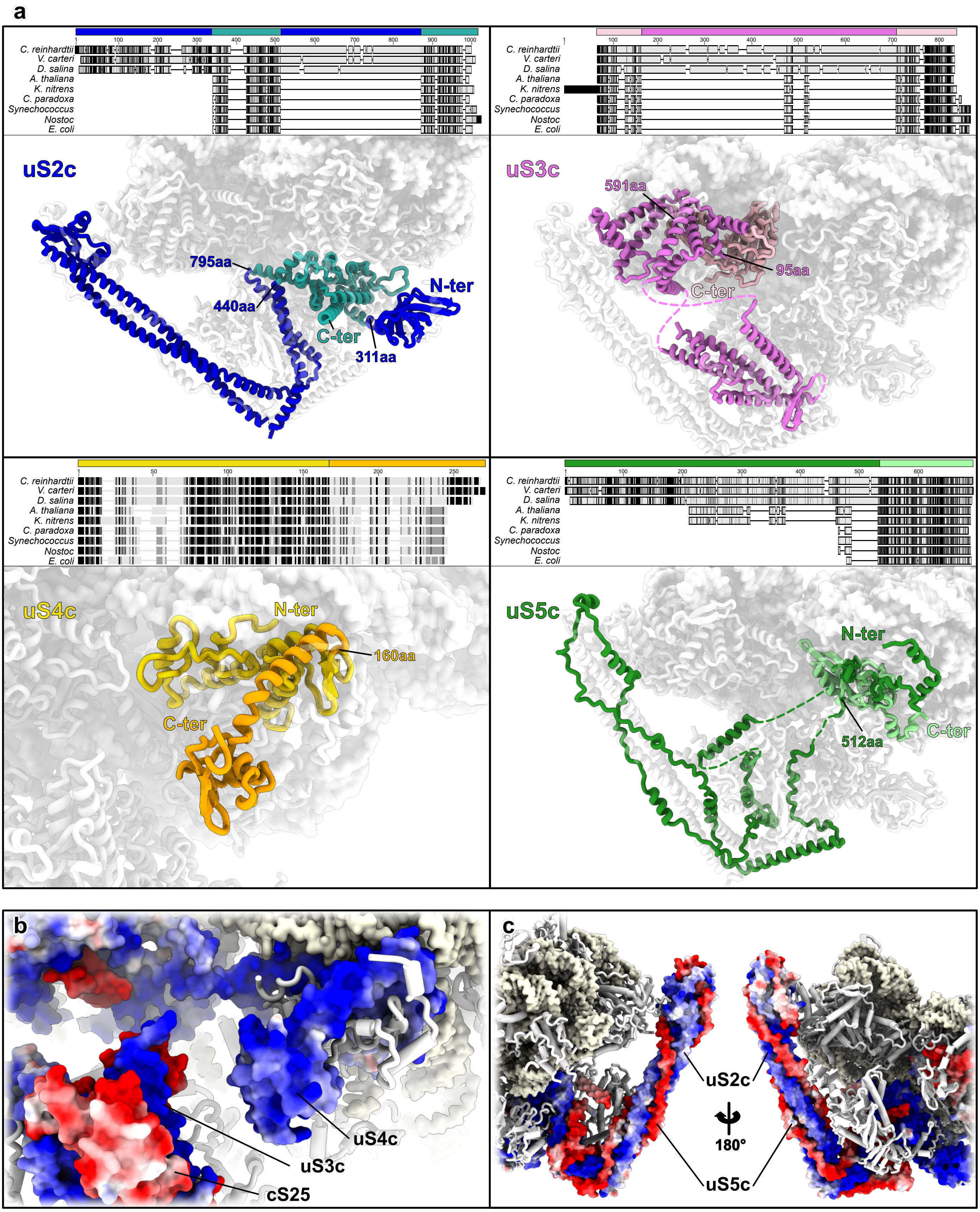


**Supplementary Figure 8:** Sequence and structural comparison of uS2c, uS3c, uS4c, and uS5c across representative taxa and charge interactions

**a** Multiple sequence alignments of uS2, uS3, uS4 and uS5 are shown for *Chlamydomonas reinhardtii*, *Volvox carteri*, and *Dunaliella salina* (Chlorophyceae); *Arabidopsis thaliana* and *Klebsormidium nitens* (Streptophyta); *Cyanophora paradoxa* (Glaucophyta); *Synechococcus elongatus* and *Nostoc sp.* PCC 7107 (Cyanobacteria); and *Escherichia coli*. Conserved core regions of each ribosomal protein are depicted in lighter shades, while algal-specific insertions and extensions are shown in darker colors, both on the 3D structures and the alignments. Unmodeled parts of the proteins are shown as dashed line. The boundaries of these extensions are annotated on the structures. uS2c (910aa) presents a 356aa insertion and a 310aa N-ter extension, uS3c (712aa) includes a 497aa insertion, uS4c (257aa) features a 97aa C-ter extension and uS5c (673aa) displays a 512aa N-ter extension. **b, c** proteins uS3c, uS4c and cS25 (**b**) as well as uS2c and uS5c (**c**) represented as surface and colored by electrostatic potential, with blue positively charged and red negatively charged.
