## Supplementary Fig. 9 for "Chloroplast-encoded small subunit extensions reshape the Chlamydomonas chlororibosome"

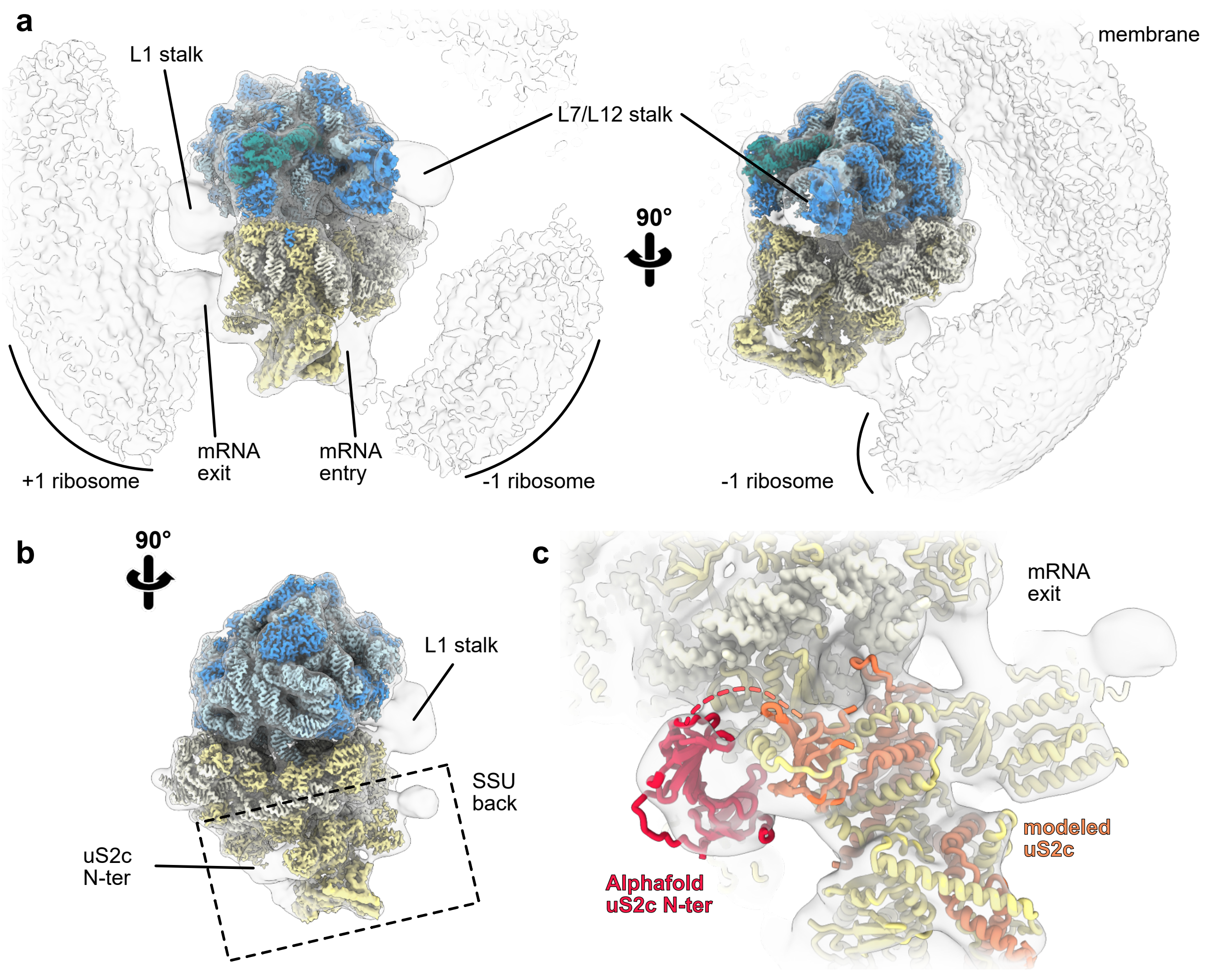


**Supplementary Figure 9:** Comparison between SPA and STA reconstructions of the chlororibosome

**a** The SPA map (colored by component: LSU rRNA, light blue; LSU ribosomal proteins, dark blue; SSU rRNA, light yellow; SSU ribosomal proteins, yellow) is shown alongside a low-pass–filtered consensus STA map (grey). Two orientations are shown, highlighting features present in STA but absent or attenuated in SPA, including additional +1 and −1 chlororibosomes within polysomes, membrane-associated density, and the L1 and L7/L12 stalk regions. **b** View of the SSU back (boxed, dashed outline), rotated 90° relative to a, revealing an unoccupied density at the expected position of the uS2c N-terminus. **c** Close-up of the SSU-back region near the mRNA exit site. The fitted chlororibosome model is shown in yellow, with the modeled uS2c segment in coral, and an AlphaFold prediction for the uS2c N-ter (residues 23–168) shown in crimson, comprising two β-barrel domains.
