## Supplementary Fig. 10 for "Chloroplast-encoded small subunit extensions reshape the Chlamydomonas chlororibosome"

## uS2c

Chlorophyceae

| Target | Description | Scientific Name | Prob. | Seq. Id. | E-Value | Position in query |
|---|---|---|---|---|---|---|
| AF-A0A0S2IDR4-F1-model_v6 | Small ribosomal subunit protein uS2c | Lobochlamys culleus | 1.00 | 43.7 | 6.92e-93 | 1–905 |
| AF-A0A2Z5DY27-F1-model_v6 | Ribosomal protein S2 | Chlamydomonas sp. UWO 241 | 1.00 | 46.8 | 1.06e-91 | 24–909 |
| AF-A0A0S2LQ45-F1-model_v6 | Small ribosomal subunit protein uS2c | Bracteacoccus giganteus | 1.00 | 40.9 | 1.54e-90 | 14–910 |
| AF-A0A1W5RMS8-F1-model_v6 | Small ribosomal subunit protein uS2c | Hydrodictyon reticulatum | 1.00 | 37.6 | 8.62e-82 | 17–908 |
| AF-A0A0S2LPR1-F1-model_v6 | Small ribosomal subunit protein uS2c | Carteria sp. SAG 8-5 | 1.00 | 44.7 | 5.28e-85 | 16–903 |
| AF-A0A0S2LMR8-F1-model_v6 | Small ribosomal subunit protein uS2c | Treubaria triappendiculata | 1.00 | 38 | 2.39e-83 | 1–909 |
| AF-A0AB39U3S7-F1-model_v6 | Small ribosomal subunit protein uS2c | Borodinellopsis insignis | 1.00 | 40.6 | 3.54e-87 | 1–904 |
| AF-A0A0S2LNK4-F1-model_v6 | Small ribosomal subunit protein uS2c | Jenufa perforata | 1.00 | 33.6 | 1.06e-76 | 1–906 |
| AF-A0A0S2IBN7-F1-model_v6 | Small ribosomal subunit protein uS2c | Microglena monadina | 1.00 | 33.9 | 7.73e-63 | 28–909 |
| AF-A0AB39A660-F1-model_v6 | Small ribosomal subunit protein uS2c | Borodinellopsis insignis | 1.00 | 53.6 | 9.92e-51 | 352–904 |
| AF-A0A0S2IBU8-F1-model_v6 | Small ribosomal subunit protein uS2c | Chlamydomonas nivalis | 1.00 | 34.5 | 4.01e-64 | 1–905 |
| AF-A0A167MGY6-F1-model_v6 | Small ribosomal subunit protein uS2c | Koshicola spirodelophila | 1.00 | 20.7 | 7.49e-17 | 302–909 |
| AF-A0A097KQK6-F1-model_v6 | Small ribosomal subunit protein uS2c | Dicloster acuatus | 1.00 | 18 | 7.75e-14 | 315–908 |

## uS3c

Chlorophyceae

| Target | Description | Scientific Name | Prob. | Seq. Id. | E-Value | Position in query |
|---|---|---|---|---|---|---|
| AF-A0A0S2LPK1-F1-model_v6 | Small ribosomal subunit protein uS3c | Chlamydomonas applanata | 1.00 | 42.2 | 8.40e-55 | 1–712 |
| AF-A0A6C0RVW8-F1-model_v6 | Small ribosomal subunit protein uS3c | Volvulina compacta | 1.00 | 62.2 | 3.87e-62 | 1–712 |
| AF-A0A5C2FSQ2-F1-model_v6 | Small ribosomal subunit protein uS3c | Chlorosarcinopsis eremi | 1.00 | 40.1 | 1.31e-50 | 1–712 |
| AF-A0A0S2IBM5-F1-model_v6 | Small ribosomal subunit protein uS3c | Lobochlamys segnis | 1.00 | 39.8 | 1.23e-51 | 1–712 |
| AF-A0A140HAA3-F1-model_v6 | Small ribosomal subunit protein uS3c | Chlorotetraedron incus | 1.00 | 37.2 | 9.21e-48 | 1–711 |
| AF-A0A0S2ICE5-F1-model_v6 | Small ribosomal subunit protein uS3c | Chloromonas radiata | 1.00 | 35.9 | 3.98e-48 | 1–712 |
| AF-B2X2C4-F1-model_v6 | Small ribosomal subunit protein uS3c | Chlamydomonas moewusii | 1.00 | 35.3 | 6.05e-48 | 1–712 |
| AF-A0A0S2LPG7-F1-model_v6 | Small ribosomal subunit protein uS3c | Carteria sp. SAG 8-5 | 1.00 | 41 | 1.65e-46 | 1–711 |
| AF-A0A2K9YS16-F1-model_v6 | Small ribosomal subunit protein uS3c | Haematococcus lacustris | 1.00 | 31.3 | 7.63e-44 | 1–712 |
| AF-A0A499SA63-F1-model_v6 | Small ribosomal subunit protein uS3c | Spermatozopsis similis | 1.00 | 37.9 | 1.28e-44 | 1–712 |
| AF-A0A0S2LNH6-F1-model_v6 | Small ribosomal subunit protein uS3c | Jenufa minuta | 1.00 | 33.8 | 7.68e-43 | 1–711 |
| AF-A0A0S2IBF7-F1-model_v6 | Small ribosomal subunit protein uS3c | Chlamydomonas nivalis | 1.00 | 32.7 | 5.87e-44 | 1–711 |
| AF-A0A140HAW2-F1-model_v6 | Small ribosomal subunit protein uS3c | Ankyra judayi | 1.00 | 42.6 | 1.42e-45 | 1–712 |
| AF-A0A0S2LMZ0-F1-model_v6 | Small ribosomal subunit protein uS3c | Oogamochlamys gigantea | 1.00 | 39.5 | 9.26e-47 | 1–711 |
| AF-A0A0S2ID77-F1-model_v6 | Small ribosomal subunit protein uS3c | Stephanosphaera pluvialis | 1.00 | 26.9 | 3.09e-37 | 1–712 |
| AF-A0A0S2LPR5-F1-model_v6 | Small ribosomal subunit protein uS3c | Staurocarteria cerasiformis | 1.00 | 33 | 6.01e-40 | 1–711 |
| AF-A0AA49LPD0-F1-model_v6 | Small ribosomal subunit protein uS3c | Johansenicoccus eremophilus | 1.00 | 31.8 | 7.15e-37 | 1–712 |
| AF-A0A1G4NU69-F1-model_v6 | Small ribosomal subunit protein uS3c | Hommersandiophycus borowitzkae | 1.00 | 51.8 | 3.70e-14 | 575–711 |
| AF-A0A3G1IWH2-F1-model_v6 | Small ribosomal subunit protein uS3c | Cyanoptyche gloeocystis | 1.00 | 48.8 | 2.32e-13 | 577–711 |
| AF-A0A3G1IW93-F1-model_v6 | Small ribosomal subunit protein uS3c | Gloeochaete wittrockiana | 1.00 | 44.3 | 1.00e-13 | 580–712 |
| AF-A0AAD8XRB5-F1-model_v6 | Small ribosomal subunit protein uS3c | Skeletonema marinoi | 1.00 | 46.6 | 3.72e-13 | 577–711 |

**Supplementary Figure 10:** Conservation of extra SSU domain (FoldSeek)

Foldseek was used to query the predicted structures of *C. reinhardtii* uS2c (top) and uS3c (bottom). Only uS2 and uS3 are shown because they presented clearly folded regions as opposed to the extensions of the other proteins that are more disordered. For each query, the table reports the top-ranked hits with the corresponding organism, Foldseek probability score, sequence identity, and E-value. This illustrates the conservation of uS2c and uS3c in chlorophyceae.
