## Supplementary Table 1 for "Chloroplast-encoded small subunit extensions reshape the Chlamydomonas chlororibosome"

| | Full high res. (PDB 9TVU - EMD-56352) | P-site tRNA (PDB 28JW - EMD-56567) | pY factor (PDB 28LU - EMD-56602) |
|---|---|---|---|
| EMDB | Unfocused EMD-56337 | Unfocused EMD-56561 | Unfocused EMD-56560 |
| **Data collection and processing** | | | |
| Magnification | 75,000x | 75,000x | 75,000x |
| Voltage (kV) | 300 | 300 | 300 |
| Electron exposure (e–/Å$^2$) | 50 | 50 | 50 |
| Defocus range (μm) | -0.8 to -2.0 | -0.8 to -2.0 | -0.8 to -2.0 |
| Pixel size (Å) | 0.84 | 0.84 | 0.84 |
| Symmetry imposed | C1 | C1 | C1 |
| Initial particle images (no.) | 1,739,330 | 1,739,330 | 1,739,330 |
| Final particle images (no.) | 225,199 | 49,141 | 175,058 |
| Map resolution (Å) - FSC 0.143 | 2.52 | 2.84 | 2.56 |
| Map resolution range (Å) | 2.3 - 5.8 | 2.4 - 7.5 | 2.3 - 6.2 |
| **Refinement** | | | |
| Initial model used (PDB code) | AF2 | AF2 | AF2 |
| FSC model (0.143) | 2.5 | 2.8 | 2.6 |
| CC Model vs. Data (mask) | 0.83 | 0.82 | 0.82 |
| Map sharpening *B* factor (Å$^2$) | -54.7 | -32.9 | -51.9 |
| Model composition | | | |
| Non-hydrogen atoms | 158411 | 160218 | 159564 |
| Residues: Protein - Nucleotide | 8399 - 4285 | 8406 - 4367 | 8545 - 4285 |
| Water | - | - | - |
| Ligands | K: 138<br>MG: 323 | K: 130<br>MG: 319<br>CLM:1 | K: 138<br>MG: 322 |
| *B* factors (Å$^2$) | | | |
| Protein | 0.25/422.72/60.49 | 1.47/442.59/85.12 | 0.00/312.71/68.79 |
| Nucleotide | 0.42/231.80/32.59 | 0.22/240.78/59.65 | 1.18/185.65/43.56 |
| Ligand | 6.73/133.09/31.00 | 20.56/119.88/52.06 | 7.9/138.32/38.73 |
| R.m.s. deviations | | | |
| Bond lengths (Å) | 0.009 | 0.007 | 0.007 |
| Bond angles (°) | 0.848 | 0.802 | 0.771 |
| Validation | | | |
| MolProbity score | 1.68 | 1.39 | 1.64 |
| Clashscore | 5.35 | 4.77 | 4.41 |
| Poor rotamers (%) | 1.77 | 2.05 | 2.20 |
| Ramachandran plot | | | |
| Favored (%) | 96.76 | 96.76 | 97.11 |
| Allowed (%) | 3.11 | 3.11 | 2.78 |
| Disallowed (%) | 0.13 | 0.13 | 0.11 |

**Supplementary Table 1:** Cryo-EM statistics table
