## Supplementary Table 2 for "Chloroplast-encoded small subunit extensions reshape the Chlamydomonas chlororibosome"

| Name | UNIPROT ID | Genome | Size | Location | Chain name | Comment |
|---|---|---|---|---|---|---|
| bS1c | Q70DX8 | Cre09.g394750 | 436 | back | a | |
| uS2c | A0A218N8X3 | CreCp.g802299 | 910 | back | b | SSU arm extension |
| uS3c | Q08365 | CreCp.g802310 | 712 | head | c | SSU arm extension |
| uS4c | P48270 | CreCp.g802283 | 257 | body | d | SSU arm extension |
| uS5c | A8J8M5 | Cre16.g659950 | 673 | body | e | SSU arm extension |
| bS6c | A8J5Y7 | Cre12.g520600 | 171 | body | f | |
| uS7c | P48267 | CreCp.g802288 | 168 | head | g | |
| uS8c | P59775 | CreCp.g802279 | 141 | body | h | |
| uS9c | O20029 | CreCp.g802303 | 191 | head | i | |
| uS10c | A8IWQ1 | Cre03.g195650 | 169 | head | j | N-ter contacts uS2c and uS5c ext. |
| uS11c | Q7YKX3 | CreCp.g802319 | 130 | body | k | |
| uS12c | P14149 | CreCp.g802325 | 133 | body | l | |
| uS13c | A8JDP6 | Cre12.g493950 | 164 | head | m | |
| uS14c | Q9GGE2 | CreCp.g802289 | 100 | head | n | |
| uS15c | A8HWZ8 | Cre06.g264300 | 141 | body | o | |
| bS16c | A8JDN8 | Cre12.g494450 | 128 | body | p | |
| uS17c | A8JGS2 | Cre02.g118950 | 105 | body | q | |
| bS18c | O20032 | CreCp.g802300 | 137 | body | r | |
| uS19c | P59776 | CreCp.g802275 | 92 | head | s | |
| bS20c | A8JDN4 | Cre12.g494750 | 166 | body | t | |
| bS21c | A8HPN4 | Cre01.g017300 | 184 | body | u | |
| cS24/PSRP3 | A8I8A3 | Cre02.g083950 | 298 | back | v | interactions with uS2c and bS1c |
| cS25/PSRP7 | Q5QEB1 | Cre12.g519180 | 560 | head | w | interactions with uS3c extensions |
| cS26/PSRP8 | A8I8A6 | Cre02.g084000 | 120 | body | x | replaces C-ter of uS4c |
| pY/PSRP1 | A8J8B2 | Cre05.g237450 | 285 | | y | only in hibernation |
| uL1c | A8I8Z4 | Cre02.g088900 | - | L1 stalk | A | not resolved |
| uL2c | Q8HTL2 | CreCp.g802274 | 278 | | B | |
| uL3c | A8JE35 | Cre48.g761197 | 259 | | C | |
| uL4c | Q84U22 | Cre11.g479500 | 243 | | D | |
| uL5c | Q8HTL1 | CreCp.g802278 | 179 | CP | E | |
| uL6c | A8J503 | Cre09.g415950 | 207 | | F | |
| bL9c | A0A2K3D5T7 | Cre12.g556050 | 200 | | G | |
| uL10c | A8HVP7 | Cre06.g272850 | 235 | L7/L12 stalk | H | |
| uL11c | A8ICE4 | Cre10.g423650 | 176 | L7/L12 stalk | I | |
| uL13c | A8HWZ6 | Cre06.g264350 | 225 | | J | |
| uL14c | P11094 | CreCp.g802277 | 122 | | K | |
| uL15c | A8JAL6 | Cre14.g612450 | 241 | | L | |
| uL16c | P05726 | CreCp.g802276 | 136 | | M | |
| bL17c | A8I3M4 | Cre02.g108850 | 173 | | N | |
| uL18c | A8HNJ8 | Cre01.g052100 | 145 | CP | O | |
| bL19c | A8IW44 | Cre17.g734450 | 153 | | P | |
| bL20c | P26565 | CreCp.g802268 | 112 | | Q | |
| bL21c | A8IP00 | Cre06.g299000 | 179 | | R | |
| uL22c | Q84U21 | Cre13.g580850 ? | 175 | | S | |
| uL23c | Q8HTL3 | CreCp.g802273 | 95 | | T | |
| uL24c | A8J9D9 | Cre16.g652550 | 170 | | U | |
| bL27c | A8INR7 | Cre06.g300800 | 161 | | V | |
| bL28c | A8HWS8 | Cre06.g265800 | 195 | | W | |
| uL29c | A8HXM1 | Cre06.g259850 | 134 | | X | |
| bL31c | A8J3Z3 | Cre08.g365400 | 136 | CP | Y | |
| bL32c | A8IUC3 | Cre07.g352850 | 98 | | Z | |
| bL33c | A8I1D3 | Cre10.g462950 | 101 | | 6 | |
| bL34c | A8HQG3 | Cre01.g030050 | 124 | | 7 | |
| bL35c | A8JEP1 | Cre04.g217932 | 114 | | 8 | |
| bL36c | P59774 | CreCp.g802272 | 37 | | 9 | |
| cL38/PSRP6 | A8IMN3 | Cre06.g308533 | 66 | | 0 | |

**Supplementary Table 2:** Ribosomal proteins of the *Chlamydomonas reinhardtii* chloroplast ribosome

Proteins are grouped by subunit (SSU, yellow; LSU, blue); chloroplast-specific proteins are highlighted in red. For each protein, the UniProt accession, gene model identifier (C.reinhardtii_CC-4532 v6.1 genome, with green indicating chloroplast encoded proteins), protein length (aa), assigned structural location, chain identifier used in the model, and relevant notes are listed.
