## Supplementary Table 3 for "Chloroplast-encoded small subunit extensions reshape the Chlamydomonas chlororibosome"

| | |
|---|---|
| tomo11 | gain1 |
| tomo12 | gain1 |
| tomo23 | gain1 |
| tomo31 | gain1 |

| | |
|---|---|
| tomo87 | gain2 |
| tomo92 | gain2 |
| tomo182 | gain2 |
| tomo200 | gain2 |
| tomo202 | gain2 |
| tomo205 | gain2 |
| tomo242 | gain2 |
| tomo243 | gain2 |
| tomo257 | gain2 |
| tomo284 | gain2 |
| tomo317 | gain2 |
| tomo318 | gain2 |
| tomo341 | gain2 |
| tomo342 | gain2 |
| tomo364 | gain2 |
| tomo373 | gain2 |
| tomo394 | gain2 |
| tomo408 | gain2 |
| tomo436 | gain2 |
| tomo444 | gain2 |
| tomo446 | gain2 |
| tomo448 | gain2 |

| | |
|---|---|
| tomo462 | gain3 |
| tomo473 | gain3 |
| tomo499 | gain3 |
| tomo517 | gain3 |
| tomo518 | gain3 |
| tomo555 | gain3 |
| tomo573 | gain3 |
| tomo576 | gain3 |
| tomo595 | gain3 |
| tomo611 | gain3 |
| tomo632 | gain3 |
| tomo663 | gain3 |
| tomo679 | gain3 |
| tomo683 | gain3 |

| | |
|---|---|
| tomo846 | gain4 |
| tomo867 | gain4 |
| tomo868 | gain4 |
| tomo869 | gain4 |
| tomo900 | gain4 |
| tomo909 | gain4 |
| tomo911 | gain4 |
| tomo912 | gain4 |
| tomo944 | gain4 |
| tomo949 | gain4 |
| tomo1034 | gain4 |
| tomo1119 | gain4 |
| tomo1132 | gain4 |
| tomo1168 | gain4 |
| tomo1182 | gain4 |
| tomo1184 | gain4 |
| tomo1198 | gain4 |
| tomo1199 | gain4 |
| tomo1212 | gain4 |
| tomo1221 | gain4 |
| tomo1272 | gain4 |
| tomo1274 | gain4 |
| tomo1283 | gain4 |
| tomo1285 | gain4 |
| tomo1286 | gain4 |
| tomo1309 | gain4 |
| tomo1310 | gain4 |
| tomo1311 | gain4 |
| tomo1315 | gain4 |
| tomo1332 | gain4 |
| tomo1334 | gain4 |
| tomo1335 | gain4 |
| tomo1341 | gain4 |
| tomo1343 | gain4 |
| tomo1345 | gain4 |

| | |
|---|---|
| tomo1402 | gain5 |
| tomo1405 | gain5 |
| tomo1533 | gain5 |

| | |
|---|---|
| tomo1711 | gain6 |
| tomo1728 | gain6 |
| tomo1743 | gain6 |
| tomo1836 | gain6 |
| tomo1855 | gain6 |
| tomo1868 | gain6 |
| tomo2325 | gain6 |
| tomo2358 | gain6 |
| tomo2362 | gain6 |
| tomo2449 | gain6 |
| tomo2453 | gain6 |
| tomo2666 | gain6 |
| tomo2668 | gain6 |
| tomo2669 | gain6 |
| tomo2680 | gain6 |
| tomo2689 | gain6 |
| tomo2730 | gain6 |
| tomo2787 | gain6 |
| tomo2824 | gain6 |
| tomo2828 | gain6 |

| | |
|---|---|
| tomo2068 | gain7 |
| tomo2122 | gain7 |
| tomo2148 | gain7 |
| tomo2152 | gain7 |
| tomo2154 | gain7 |
| tomo2164 | gain7 |
| tomo2183 | gain7 |
| tomo2190 | gain7 |
| tomo2196 | gain7 |
| tomo2203 | gain7 |
| tomo2204 | gain7 |
| tomo2212 | gain7 |
| tomo2245 | gain7 |
| tomo2271 | gain7 |
| tomo2278 | gain7 |
| tomo2283 | gain7 |
| tomo2285 | gain7 |

| | |
|---|---|
| tomo2839 | gain7_2° |
| tomo2871 | gain7_2° |
| tomo2879 | gain7_2° |
| tomo2882 | gain7_2° |
| tomo2885 | gain7_2° |
| tomo2893 | gain7_2° |
| tomo2895 | gain7_2° |
| tomo2902 | gain7_2° |
| tomo2904 | gain7_2° |
| tomo2905 | gain7_2° |
| tomo2908 | gain7_2° |
| tomo2914 | gain7_2° |
| tomo2942 | gain7_2° |
| tomo2946 | gain7_2° |

Gain references:

| | |
|---|---|
| gain1 | 20211216_124009_EER_GainReference.gain |
| gain2 | 20220427_000718_EER_GainReference.gain |
| gain3 | 20220717_000157_EER_GainReference.gain |
| gain4 | 20220717_000157_EER_GainReference.gain |
| gain5 | 20230807_182744_EER_GainReference.gain |
| gain6 | 20230811_112217_EER_GainReference.gain |
| gain7 | 20230903_185905_EER_GainReference_NNPKrios.gain |

**Supplementary Table 3:** List of tomograms used from EMPIAR-11830

List of the tomograms used from the publicly deposited EMPIAR-11830. The tomograms are broken down by gain reference, and the last ones were collected with a 2° tilt increment, contrary to the rest that were collected with a 3° increment. The correspondance between the tomogram numbers and their raw names in the deposited can be found at https://github.com/Chromatin-Structure-Rhythms-Lab/ChlamyAnnotations.
